# Overnutrition directly impairs thyroid hormone biosynthesis and utilization, causing hypothyroidism, despite remarkable thyroidal adaptations

**DOI:** 10.1101/2025.03.31.645596

**Authors:** Jessica Rampy, Alejandra Paola Torres-Manzo, Kendra Hoffsmith, Matthew A. Loberg, Quanhu Sheng, Federico Salas-Lucia, Antonio C. Bianco, Rafael Arrojo e Drigo, Huiying Wang, Vivian L. Weiss, Nancy Carrasco

**Affiliations:** Department of Cellular & Molecular Physiology, Yale University, New Haven, CT; Department of Molecular Physiology & Biophysics, Vanderbilt University, Nashville, TN; Department of Pathology, Microbiology, & Immunology, Vanderbilt University Medical Center, Nashville, TN; Department of Biostatistics, Vanderbilt University Medical Center, Nashville, TN; Department of Medicine, The University of Chicago, Chicago, IL; Center for Computational Systems Biology, Vanderbilt University, Nashville, TN

## Abstract

Thyroid hormones (THs: T_3_ and T_4_) are key regulators of metabolic rate and nutrient metabolism. They are controlled centrally and peripherally in a coordinated manner to elegantly match T_3_-mediated energy expenditure (EE) to energy availability. Hypothyroidism reduces EE and has long been blamed for obesity; however, emerging evidence suggests that, instead, obesity may drive thyroid dysfunction. Thus, we used a mouse model of diet-induced obesity to determine its direct effects on thyroid histopathology and function, deiodinase activity, and T_3_ action. Strikingly, overnutrition induced hypothyroidism within 3 weeks. Levels of thyroidal THs and their precursor protein thyroglobulin decreased, and ER stress was induced, indicating that thyroid function was directly impaired. We also observed pronounced histological and vascular expansion in the thyroid. Overnutrition additionally suppressed T_4_ activation, rendering the mice resistant to T_4_ and reducing EE. Our findings collectively show that overnutrition deals a double strike to TH biosynthesis and action, despite large efforts to adapt—but, fortunately, thyroid dysfunction in mice can be reversed by weight loss. In humans, BMI correlated with thyroidal vascularization, importantly demonstrating preliminary translatability. These studies lay the groundwork for novel obesity therapies that tackle hypothyroidism—which are much-needed, as no current obesity treatment works for everyone.

## Introduction

The thyroid hormones (THs: T_3_ and T_4_) are required for physiological regulation of energy homeostasis, carbohydrate and lipid metabolism, thermogenesis, and cellular metabolism (1). Thyroid function is directly regulated by thyroid stimulating hormone (TSH), and THs in turn suppress TSH synthesis and release by the pituitary. This negative feedback loop is part of the hypothalamic–pituitary–thyroid (HPT) axis. When TH levels are low, TSH levels rise to stimulate every major step of TH biosynthesis (2). Thyrocytes synthesize and properly fold the ∼660 kDa protein dimer thyroglobulin (TG) and secrete it into the follicular colloid. Iodide (I^-^) is actively transported by the Na^+^/I^-^ symporter (NIS) from the blood into the thyrocytes (3–5) and eventually gets incorporated onto tyrosyl residues of TG in the colloid. Finally, iodinated TG is endocytosed back into the thyrocytes, where the THs are cleaved from TG and released into the blood.

It has been known for over a century that THs are potent stimulators of energy expenditure (EE) and that hypothyroidism reduces EE (6). In fact, basal metabolic rate (BMR) is directly correlated with serum T_4_ levels (7). Hypothyroidism, even when mild, also causes other metabolic dysfunctions, including hyperlipidemia, cardiovascular disease, and metabolic dysfunction–associated steatotic liver disease (MASLD) (8–12). Whether hypothyroidism causes weight gain is intensely debated: several studies report an inverse association between thyroid function and BMI (13–15), whereas others demonstrate that treatment of hypothyroidism results in only modest weight loss, attributable to appropriate fluid loss in some cases (16, 17). Even so, obesity is, on the whole, associated with normal or high TSH levels and normal or low free T_4_ levels (16), as well as with an increased prevalence of hypothyroidism (18). Due to THs’ potent effects on EE, it has long been postulated that obesity is secondary to hypothyroidism when these disorders coincide. However, the lack of compelling data linking primary hypothyroidism to weight gain calls this assumption into question. Rather, mounting evidence suggests that obesity can induce thyroid dysfunction, in line with its deleterious effects on many other hormonal systems (16). For instance, obesity is associated with greater thyroid volume, attributable to elevated TSH (19), and with a hypoechogenic ultrasound pattern in the thyroid, a diagnostic sign of reduced thyroid function (20, 21). Moreover, a meta-analysis of bariatric patients revealed that preoperative TSH and serum T_3_:T_4_ were high (22). All these phenomena are routinely interpreted as evidence of inadequate thyroid function in various forms of hypothyroidism and are improved by weight loss (22–28).

It has been hypothesized that elevated TSH in obese subjects is a physiological adjustment to increase TH levels and EE to combat positive energy balance (29). However, studies show that elevation of serum T_3_ in response to overfeeding is a homeostatic mechanism for increasing EE that does not require increased TSH stimulation of the thyroid in lean subjects (30–32). Mechanistically, this pattern is accomplished by increased peripheral T_4_-to-T_3_ conversion by deiodinases (33). T_4_ is converted to T_3_ by deiodinase 1 (D1) and deiodinase 2 (D2), which are highly expressed in liver and brown adipose tissue (BAT), respectively (34). T_4_ can also be inactivated by deiodinase 3 (D3), which is abundant in the brain. The thyroid secretes mostly the prohormone T_4_; most of the body’s T_3_, the active hormone, results from D2-mediated deiodination of T_4_ (35), demonstrating that peripheral mechanisms play a key role in regulating serum TH levels. A recent meta-analysis of overfeeding in lean subjects concurred that overnutrition increased T_3_—and consequently T_3_:T_4_—in most studies, but not a single study found a change in TSH (36). In contrast, the co-occurrence of elevated TSH with increased T_3_:T_4_ in obese subjects, along with the other thyroid symptoms described, make a strong case for obesity-induced thyroid dysfunction, not simply physiological adaptation. Data regarding how obese subjects respond to experimental overfeeding is sparse, underpowered, and conflicting (37–39). Furthermore, the effect of obesity on deiodinase activity is not yet well understood but could dampen the body’s ability to upregulate EE.

To further elucidate the effects of obesity on thyroid function and TH action, we used a mouse model of overnutrition to interrogate its effects on systemic TH levels and thyroid histology and function, with a particular emphasis on identifying biochemical mechanisms by which short-term overnutrition may impair thyrocyte activity. Our results indicate that overnutrition directly impairs thyroid function while also triggering remarkable, albeit insufficient, compensatory mechanisms aimed at increasing TH biosynthesis within the thyroid—a phenotype mostly reversed by weight loss. We also found that overnutrition modulates deiodinase activity, reducing whole-body utilization of T_4_ and EE. Our preliminary study in human thyroids revealed similar histological changes associated with high BMI. These findings collectively support the hypothesis that thyroid dysfunction may be secondary to obesity in many patients, and highlight the key role that TH action plays in increasing metabolic rate. Furthermore, TH action could be harnessed to promote weight loss that could ultimately repair obesity-related thyroid damage.

## Results

### Overnutrition induces hypothyroidism and goiter

Male C57Bl/6J mice were placed on a high-fat food plus sucrose water diet (HF+SD) for a short period of time (6 weeks; Figure 1A). These mice consumed more calories than control mice and increased their BW and fat mass (Figure 1, B–D). Whereas serum T_3_ levels were maintained, serum T_4_ decreased and TSH increased within 3 weeks, the latter worsening in a time-dependent manner (Figure 1, E–G). The HF+SD-fed mice also progressively developed goiter within 3 weeks (Figure 1H). Goiter is a classic symptom of hypothyroidism resulting from a sustained elevation in TSH levels, which is well known to have mitogenic effects (2). These results indicate that overnutrition induces mild hypothyroidism in this mouse model. The diet-induced hypothyroidism was not due to I^-^ deficiency, as I^-^ excretion was not different between groups at any timepoint, nor was the elevated TSH attributable to stimulation by leptin, because leptin was unchanged between groups (Supplemental Figure 1). HF+SD-fed female mice exhibited a similar, albeit milder, systemic thyroid status, whereas their goiters were as pronounced as those of their male counterparts. Interestingly, their BW increase was only a trend, and their fat mass was unchanged relative to that of the controls (Supplemental Figure 2). Thus, the diet-induced hypothyroid phenotype, particularly its goiter component, is observed in both sexes.

**Figure 1.**
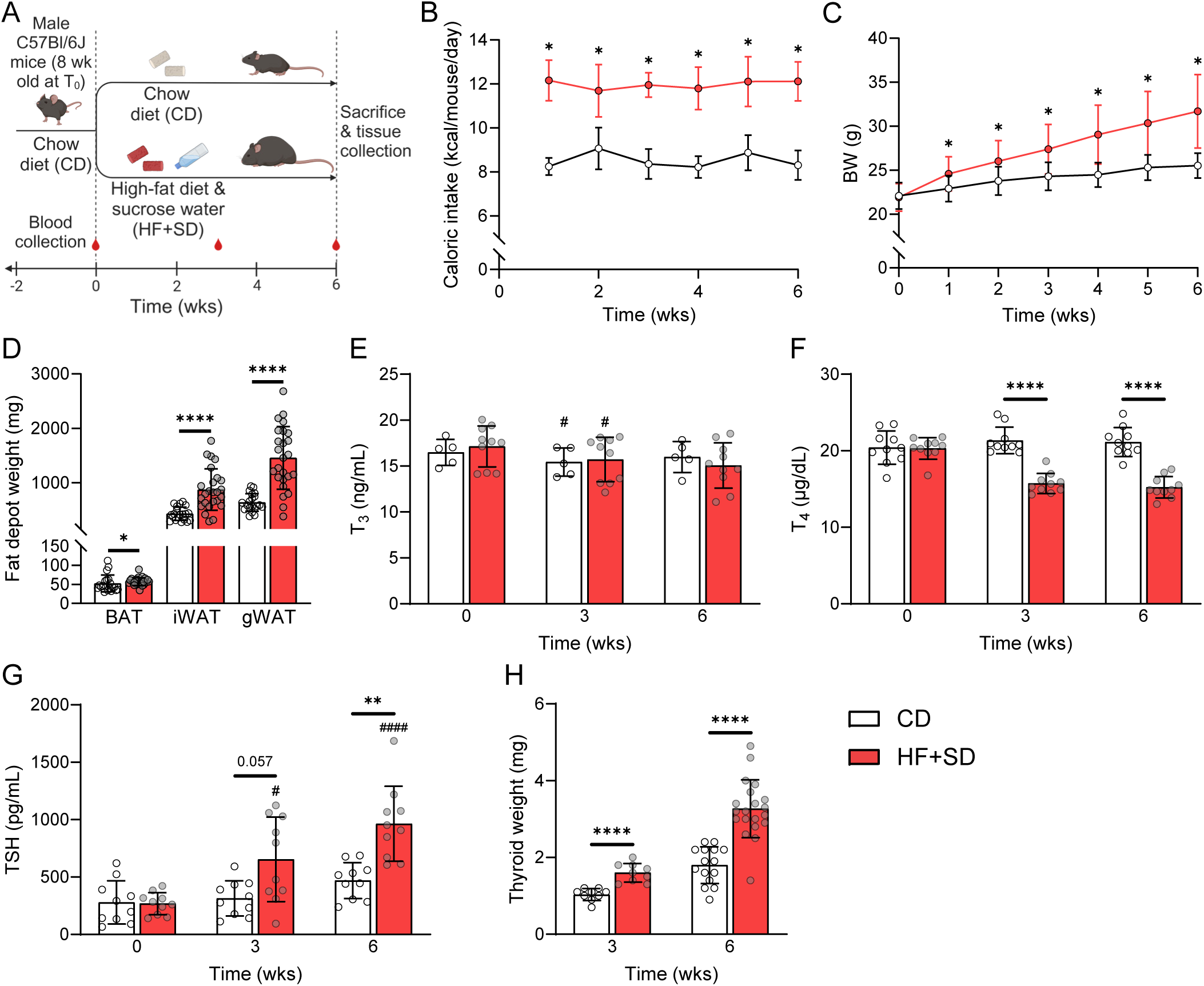
Overnutrition induces hypothyroidism and goiter. All studies followed this general design (**A**) unless otherwise specified. Caloric intake (**B**) and BW (**C**) were measured weekly. BAT, inguinal white adipose tissue (iWAT), and gonadal WAT (gWAT) depots were weighed immediately after dissection (**D**). Plasma and sera were analyzed by immunoassay for total T_3_ (**E**), total T_4_ (**F**) and TSH (**G**). Thyroids were carefully dissected at either week 3 or week 6 and weighed (**H**). All data are representative of at least two separate cohorts of mice: this cohort CD *n* = 20 in 4 cages, HF+SD *n* = 25 in 5 cages (B–D). Data were analyzed by 2-way ANOVA with repeated measures (B, C, E–G), unpaired Student’s *t*-test with Welch’s correction (D: WAT, H), or Mann-Whitney test (D: BAT). \**p* < 0.05, \*\**p* < 0.01, & \*\*\*\**p* < 0.0001 vs. controls within that timepoint. #*p* < 0.05 & ####*p* < 0.0001 vs. week 0 within that diet group.

### Overnutrition reduces T_4_-to-T_3_ conversion and EE by impairing D2 activity

To investigate TH metabolism, we measured deiodination rates in tissues from male mice on the HF+SD for 6 weeks. There was no difference in D3 activity in either the hippocampus or the cortex (Figure 2A), suggesting that D3-mediated inactivation of T_4_ is not a major contributor to diet-induced hypothyroidism. In contrast, liver D1 activity increased (Figure 2B). However, the effect of high liver D1 on serum T_4_ levels is difficult to interpret, since D1 has a comparatively poor affinity for T_4_ and catalyzes activating and deactivating deiodination of T_4_ at similar rates. Though D1 contributes minorly to the serum T_3_ pool, its primary function is to recycle I^-^ by removing it from TH degradation products (40, 41). Even so, it is possible that this increase in D1 activity lowers serum T_4_ levels. Finally, BAT D2 activity tended to be lower in the HF+SD-fed mice than in the control mice (Figure 2C), consistent with a similar trend in its protein expression (Supplemental Figure 3, A and B). This decrease in D2 activity would be expected to preserve serum T_4_ levels rather than deplete them, indicating that D2 action, at least in adipose tissue, is not a major contributor to diet-induced hypothyroxinemia. Similar results for the three deiodinases were obtained in female mice (Supplemental Figure 3, C–E).

**Figure 2.**
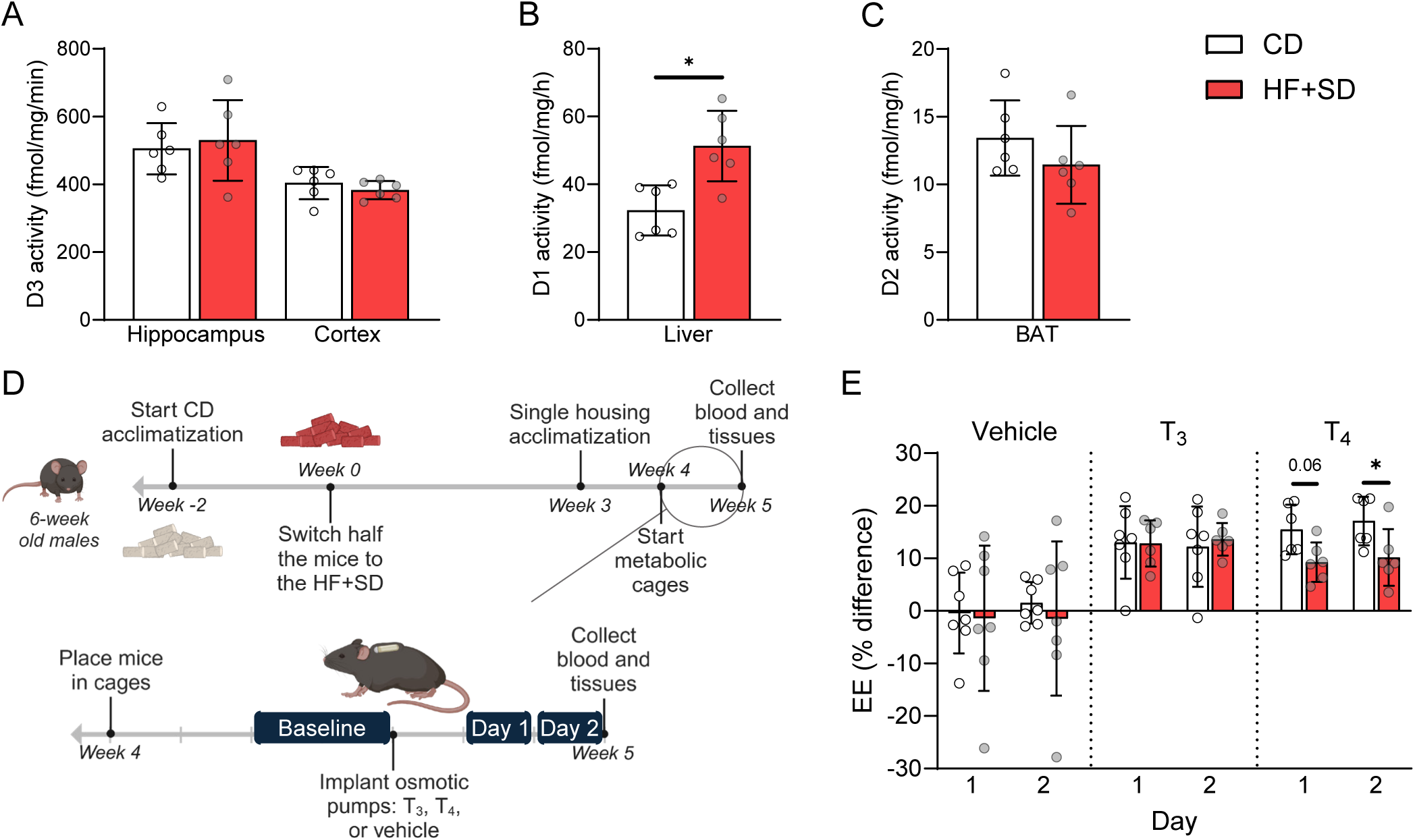
Overnutrition modulates deiodinase activity, reducing local T_4_-to-T_3_ conversion and EE overall. Deiodination rates mediated by D3 in hippocampus and cortex (**A**), D1 in liver (**B**), and D2 in BAT (**C**) were measured. The metabolic cage study design is shown (**D**). EE was measured at baseline for 48 h and then in response to vehicle, T_3_, or T_4_ for two 24-h time periods (“Day 1” and “Day 2”) after osmotic minipump implantation. The percent difference of the average EE between the response “Days” and baseline was then calculated for each mouse (**E**). Data were analyzed by unpaired Student’s *t*-test with Welch’s correction (A–C) or 2-way ANOVA with repeated measures (E). \**p* < 0.05.

To determine the combined effects of these overnutrition-induced changes in deiodinase activity on whole-body TH utilization, we performed a metabolic cage experiment quantitating the energetic response to TH administration (Figure 2D). Not surprisingly, all the mice receiving exogenous THs increased their EE (Figure 2E). The diet groups responded similarly to T_3_, suggesting that TH degradation rates are unchanged by the HF+SD, consistent with our D3 activity assay results. However, the HF+SD-fed mice exhibited a reduced response to T_4_ than did the lean mice. Furthermore, differences in EE were not due to differences in locomotor activity—or in serum TSH, T_4_, or, notably, T_3_ levels (Supplemental Figure 4). This outcome suggests that any increase in serum T_3_ levels mediated by the increased hepatic D1 activity is offset by decreased production of T_3_ elsewhere. Moreover, overnutrition impairs intracellular conversion of T_4_ to T_3_ in highly metabolic tissues to a degree that reduces overall EE. This result is most simply explained by decreased D2 activity, likely in various highly metabolic tissues, given that the differences we observed in BAT were small. Thus, it is clear that the ultimate effect of overnutrition on peripheral TH metabolism is to decrease local activation of T_4_ and slow down metabolic rate.

### Overnutrition induces hypothyroidism by directly impairing thyroid function

The reduced serum T_4_ levels caused by overnutrition could result from reduced TH biosynthesis. Thus, we characterized TH biosynthesis at various study endpoints directly by measuring the TH content in proteolytically digested thyroid homogenates; this is mostly de novo synthesized TH, since most TH in the thyroid would be covalently bound to TG prior to proteolytic digestion. Strikingly, the thyroidal T_4_ content of male mice was reduced after just 1.5 weeks of HF+SD feeding and decreased progressively (Figure 3A). Thyroidal T_3_ biosynthesis remained unchanged during the first 6 weeks on the HF+SD but eventually declined compared to controls after 12 weeks of overnutrition (Figure 3B). We also observed a robust diet-induced increase in the thyroidal T_3_:T_4_ ratio, a well characterized response to TSH stimulation and thyroid disease (42), at all timepoints (Figure 3C), in line with the notably lower efficiency of TH biosynthesis (total TH/protein content) in the HF+SD group. Thus, despite continued thyroid growth after 12 weeks of the HF+SD (Figure 3D), overnutrition induces hypothyroidism by directly impairing thyroid function.

**Figure 3.**
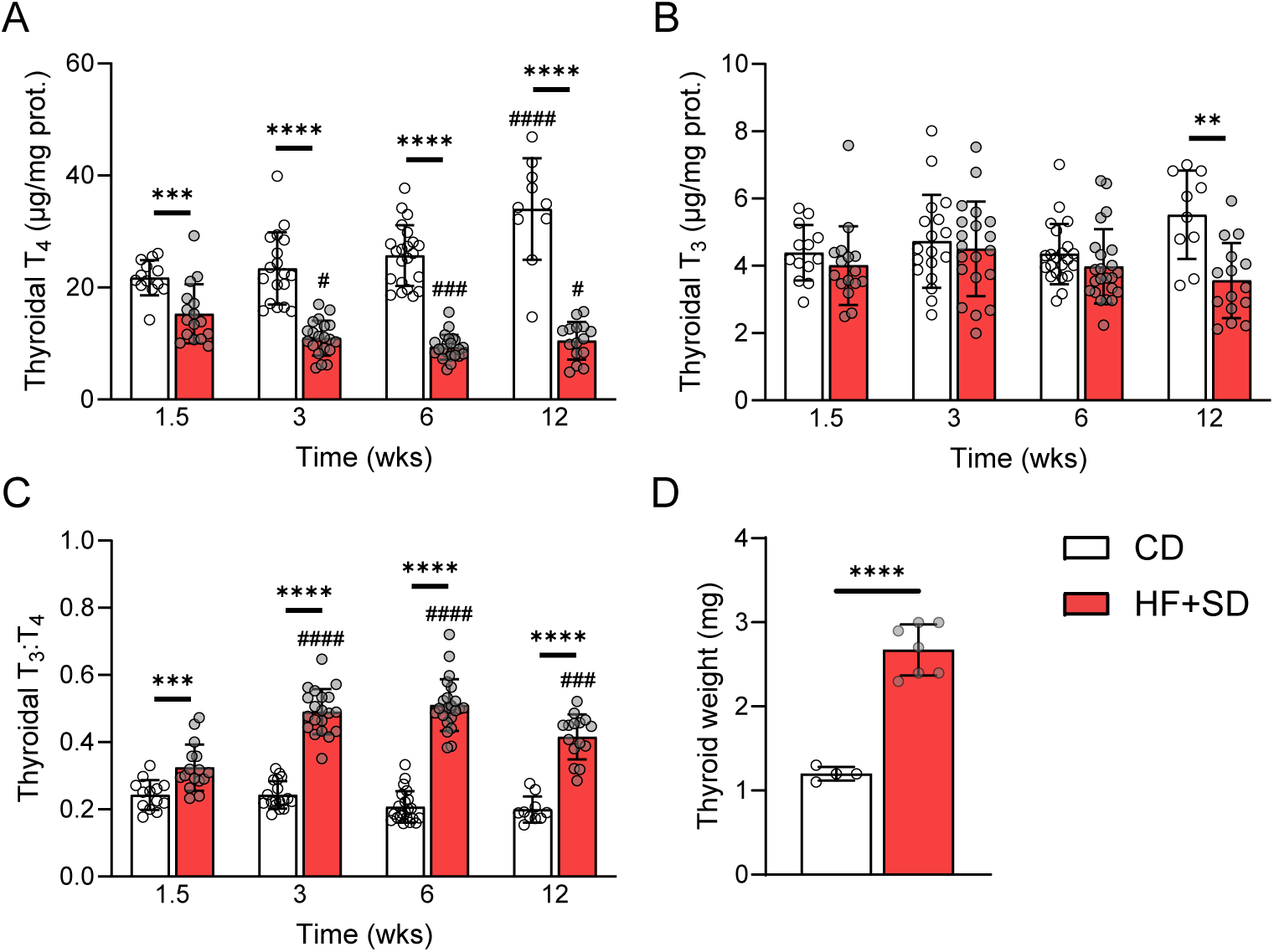
Overnutrition directly impairs TH biosynthesis. Male mice were placed on the HF+SD or CD for up to 12 weeks. At each timepoint, thyroids were collected and fully proteolyzed. Liberated T_4_ (**A**) and T_3_ (**B**) were measured by immunoassay and normalized to the thyroidal protein. The thyroidal T_3_:T_4_ ratio was calculated (**C**). At the 12-week timepoint, some thyroids were collected and weighed (**D**). Data are results from 2-3 separate cohorts of mice per timepoint: combined CD *n* = 10-22, HF+SD *n* = 15-23 (A–C). Data were analyzed per timepoint by unpaired Student’s *t*-test with Welch’s correction or Mann-Whitney test, based on normality, and across timepoints by 2-way ANOVA with multiple comparisons. \*\**p* < 0.01, \*\*\**p* < 0.001, & \*\*\*\**p* < 0.0001 vs. controls at that timepoint. #*p* < 0.05, ###*p* < 0.001, & ####*p* < 0.0001 vs. week 1.5 within that diet group.

### Overnutrition leads to pronounced histological changes in the thyroid

To identify potential mechanisms by which overnutrition directly impairs thyroid function, we characterized its effect on the histopathology of the thyroid. Healthy thyrocytes from CD-fed mice were thin and flat, with a high nuclear-to-cytoplasmic (N/C) ratio (Figure 4A). The HF+SD greatly expanded the thyrocytes’ cytoplasmic volume (thyroids with high N/C ratio: HF+SD = 0% vs. CD = 100%) and granularity, indicating greater organelle content (Figure 4, B and C). These results are consistent with the classic mitogenic effects of high TSH levels (2, 43), suggesting that TSH signaling remains at least partially intact. Furthermore, the thyroids from the HF+SD mice displayed larger interfollicular vascular/lymphatic space (Figure 4D). Since no immune cell invasion was observed, we performed immunohistochemical staining with the vascular marker CD31, revealing that overnutrition increased both the circumferential coverage of thyroid follicles by microcapillaries and their dilatation (Figure 4, E–H). We observed similar results in males after just 3 weeks and in females after 6 weeks (Supplemental Figure 5). The overall picture that emerges from these histological findings is one in which the thyroid gland adapts to overnutrition by increasing its TH biosynthesis and delivery capacity. Although this state is consistent with the effects of TSH stimulation, it does not explain the impaired thyroid function we observed.

**Figure 4.**
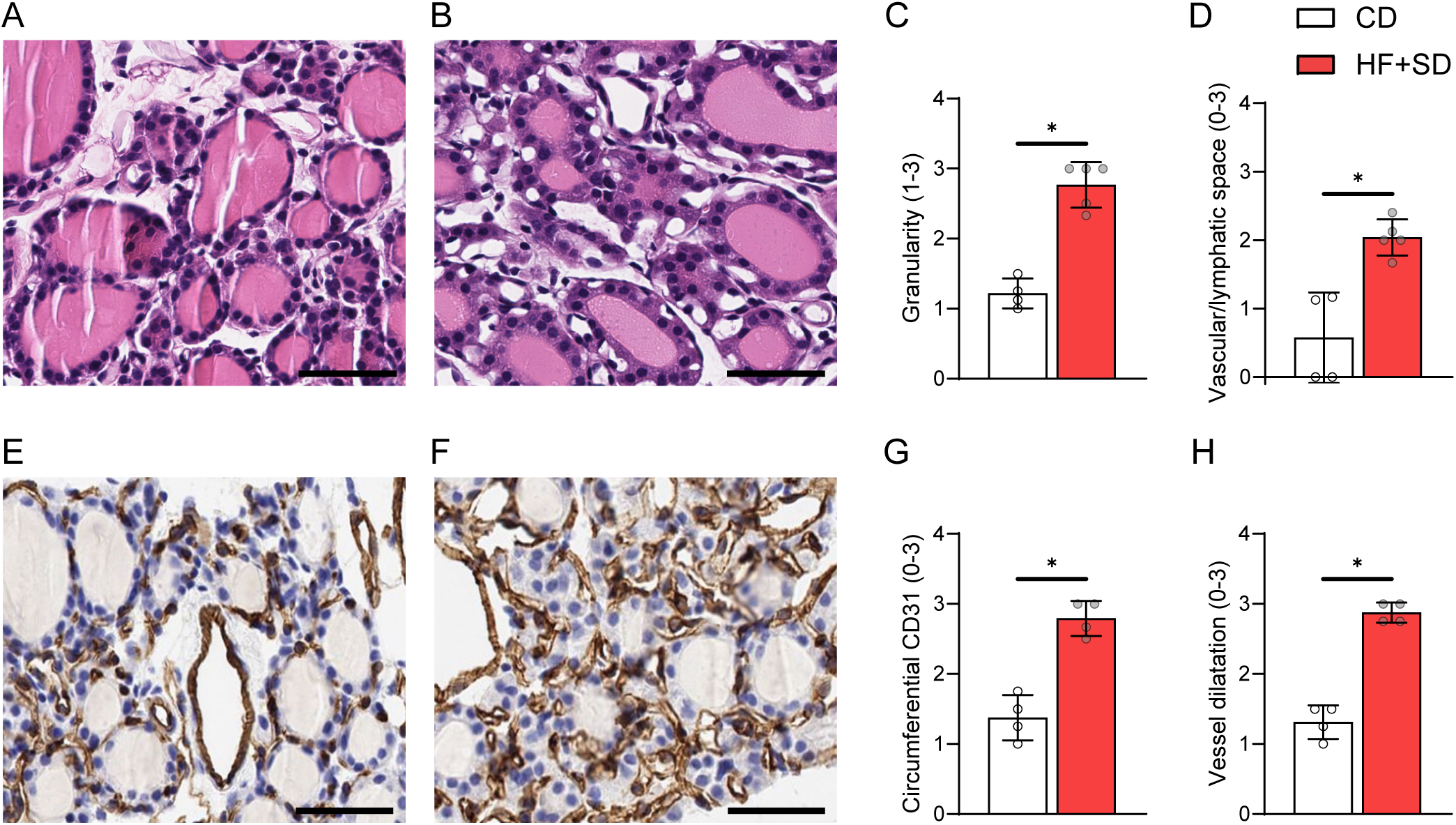
Overnutrition leads to hypertrophy, increased granularity, and increased vascularization of thyrocytes. Representative H&E images from the CD (**A**) and HF+SD (**B**) groups are shown. H&E images were blindly scored for extent of granularity (**C**) and interfollicular vascular/ lymphatic space (**D**). Fixed thyroid sections were also stained for CD31, a vascular marker, and images from CD mice (**E**) and HF+SD mice (**F**) were blindly scored for extent of circumferential staining around the follicles (**G**) and dilatation of the follicular microcapillaries (**H**). Scale bars = 50 µm. Data were analyzed by Mann-Whitney test. \**p* < 0.05.

### Overnutrition induces thyroidal ER stress, limits TG synthesis, and alters mitochondria

Next, we investigated the morphological adaptation more deeply by electron microscopy (EM). The hypertrophy of the thyrocytes, their increased granularity, and the increased and dilated follicular vasculature of the HF+SD mice were even more apparent at a higher magnification (Figure 5, A and B). We employed deep learning models to quantitate several morphological parameters, achieving 70–90% accuracy (Supplemental Figure 6, A–C). As was already apparent, 6 weeks of the HF+SD caused thyrocyte hypertrophy and increased granularity, mediated mostly by expansion of the ER and increased mitochondrial density (Figure 5, C and D; Supplemental Figure 6, D–G). However, only complex IV was upregulated, whereas other complexes were expressed at the same levels as in control mice. Moreover, complex V—the ATP synthase—was notably downregulated (Supplemental Figure 7), suggesting that overnutrition may induce some degree of mitochondrial dysfunction. If production of ATP is impaired, its availability may be a factor limiting TH biosynthesis, a possibility that should be investigated in further studies.

**Figure 5.**
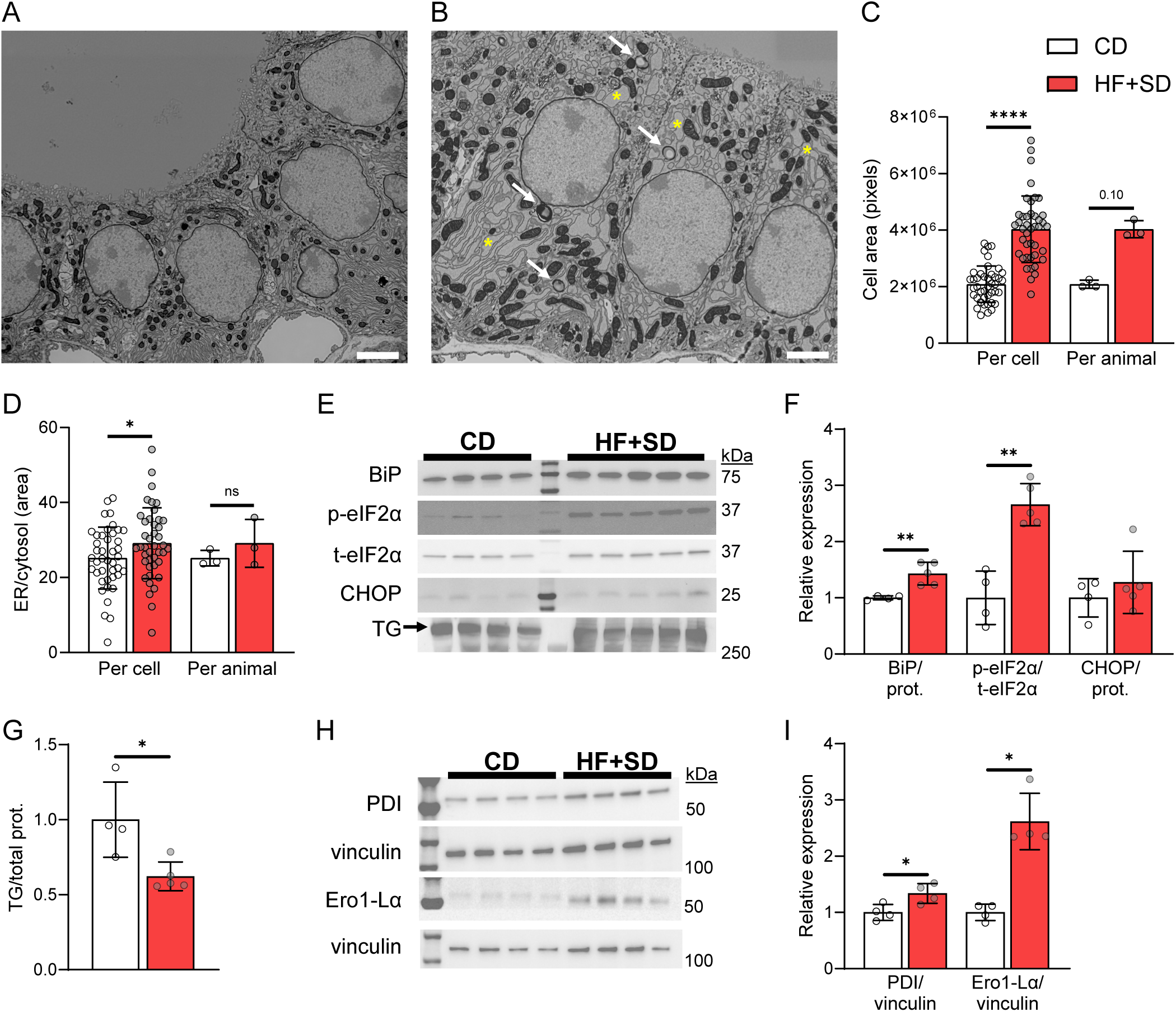
Overnutrition induces thyroidal ER stress and reduces TG. Thyroids were processed for EM. Representative images from a CD mouse (**A**) and a HF+SD mouse (**B**) are shown. Yellow asterisks label a few examples of bloated ER. White arrows point to lipid droplets. Deep learning models were used to quantitate thyrocyte total area (**C**) and ER area relative to cytosol area (**D**). Western blots of thyroid homogenate (**E** and **H**) were performed and quantitated for ER stress markers (**F**), TG (**G**), and TG-folding chaperones (**I**). Scale bars = 2.5 µm. EM data are presented per cell (*n* = 42/group) and as the average of the 14 cells/animal. Data were analyzed by unpaired Student’s *t*-test with Welch’s correction or Mann-Whitney test, based on normality. \**p* < 0.05, \*\**p* < 0.01, & \*\*\*\**p* < 0.0001.

Thyrocytes are secretory cells, producing large amounts of TG for TH biosynthesis. TG synthesis requires considerable energy and resources, as it involves the translation and folding of the ∼660 kDa dimer in the ER, post-translational modifications in the ER and Golgi, and vesicular secretion into the colloid (42). It is thus no surprise that thyrocytes contain abundant ER and that stimulation by TSH causes the ER and mitochondrial compartments to expand. However, the thyrocytes from the HF+SD mice exhibited more ER bloating than did controls, suggesting ER stress (Figure 5, A and B). We measured several markers of ER stress, revealing that overnutrition increased thyroidal expression of molecular chaperone BiP and phosphorylation of translation initiation factor eIF2α, whereas the expression of transcription factor CHOP remained unchanged (Figure 5, E and F; Supplemental Figure 8A). These results indicate ER stress severe enough to arrest global translation but not induce apoptosis (44, 45). Even so, the halting of translation could have a deleterious effect on the function of thyrocytes, as it should limit the synthesis of TG. Indeed, TG expression was decreased (Figure 5, E and G; Supplemental Figure 8B), even though *Tg* mRNA transcript levels were slightly elevated in the thyroids of HF+SD mice (Table 1). In addition, the protein expression levels of two molecular chaperones critical for proper TG folding—PDI and Ero1-Lα—were elevated (Figure 5, H and I). Thus, although the EM data are consistent with the picture of a thyroid gland adapting to increase its TH biosynthesis and delivery capacity, ER stress likely impairs TG biosynthesis and ultimately limits TH biosynthesis, resulting in thyroid dysfunction.

**Table 1.** TSH-responsive gene expression.

| Gene | Log<sub>2</sub> FC | p-value |
| --- | --- | --- |
| *Slc5a5* (NIS) | 3.1 | 1.4E-18 |
| *Tshr* | 1.7 | 1.2E-07 |
| *Tpo* | 2.3 | 4.7E-11 |
| *Duox2* | 0.8 | 0.001 |
| *Tg* | 0.6 | 0.01 |
| *Dio1* | 2.0 | 9.3E-17 |

Thyroids were processed for RNA sequencing, and differential gene expression analysis was performed. The log<sub>2</sub> fold change (FC) of TSH-responsive genes expressed in the HF+SD group vs. controls are shown, alongside adjusted *p*-values. CD *n* = 3, HF+SD *n* = 4.

Since lipotoxicity induces ER stress (46–48), we measured total triglycerides and an array of sphingolipid species in thyroid tissue, revealing mostly similar and even decreased lipid levels in the HF+SD group (Supplemental Figure 9). Thus, ceramide accumulation does not account for the thyrocyte dysfunction we observed. In fact, increased dihydroxy-sphingosine without an increase in downstream ceramides, as we observed, can itself result from ER stress (49). Only EM was sensitive enough to reveal lipid deposition (Figure 5B), and quantitative analysis showed a shift in the percentage of thyrocytes containing one or more lipid droplets (LDs) in the HF+SD group (1 LD: HF+SD = 26.2% vs. CD = 9.5%; >1 LD: HF+SD = 26.2% vs. CD = 4.8%; *n* = 42 cells/group). However, almost half the HF+SD thyrocytes did not contain any LDs, and it is unclear whether the small number of LDs observed could induce such marked ER stress and thyrocyte dysfunction.

### High TSH signaling appears unimpaired and likely drives the increased thyroidal vascularization by upregulating adrenomedullin 2 (ADM2) expression in thyrocytes

We performed bulk RNA sequencing of mouse thyroid tissue to investigate other possible mechanisms underlying the diet-induced thyroid phenotype. Six weeks of the HF+SD induced differential expression of over 6,000 genes (Figure 6A). Gene set enrichment analysis (GSEA) using gene ontology (GO) annotations yielded many top upregulated gene sets related to cellular proliferation (Supplemental Table 1), suggesting that the observed goiter was due to hyperplasia in addition to hypertrophy, consistent with the mitogenic effects of TSH (Figure 1, G and H; Figure 5C). The most downregulated gene sets were related to mitochondrial structure and function and translation, in line with our previous findings suggesting changes in mitochondria and ER stress (Supplemental Figures 6 and 7; Figure 5). Additional downregulated translation-related gene sets appeared after 3 weeks of overnutrition, suggesting that overnutrition may induce ER stress quite rapidly in the thyroid (Supplemental Table 2).

**Figure 6.**
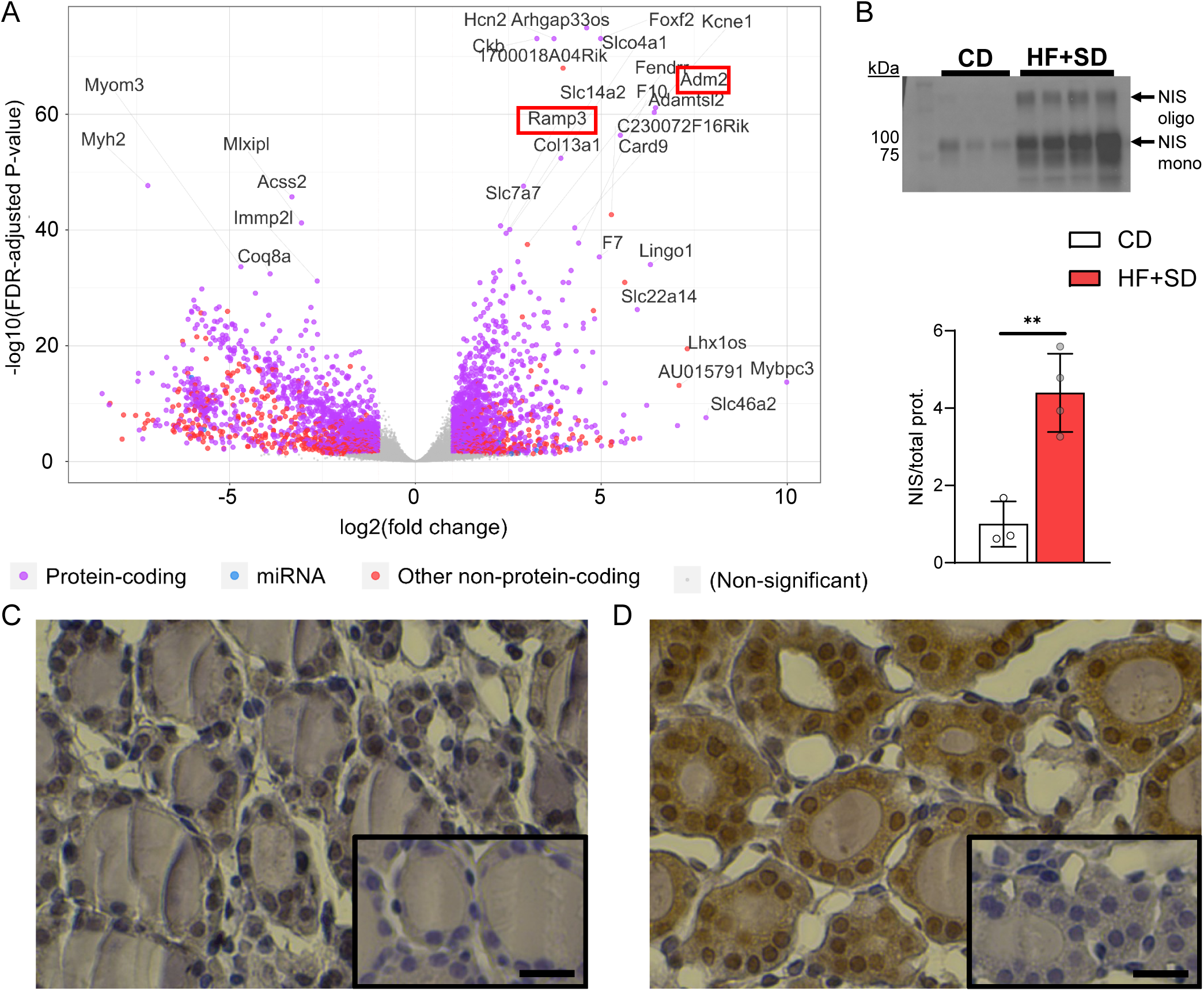
High TSH signaling appears unimpaired and likely drives thyroidal vascularization by upregulating thyrocyte ADM2 expression. Thyroids were processed for RNA sequencing. Differential gene expression analysis yielded over 6,000 genes (**A**). Western blot of thyroid homogenate was performed for NIS and quantitated (**B**). The lower arrow indicates fully glycosylated monomeric NIS. The upper arrow indicates oligomeric NIS, and the other bands correspond to partially glycosylated NIS. Fixed thyroid sections were stained for ADM2, and representative images from the CD (**C**) and HF+SD (**D**) groups are shown (insets: negative controls lacking primary antibody). Scale bars = 20 µm. CD *n* = 3, HF+SD *n* = 4, log_2_ FC > 1, & FDR-adjusted *p*-value < 0.05 (A). Data were analyzed by unpaired Student’s *t*-test with Welch’s correction (B). \*\**p* < 0.01.

Another hypothesis is that perhaps the modest ectopic lipid deposition observed was enough to induce resistance to the elevated TSH, akin to overnutrition-induced insulin and leptin resistance (16). However, NIS protein expression was highly upregulated in both male and female mice on the HF+SD (Figure 6B; Supplemental Figure 10), and TSH-responsive genes were expressed at higher levels in the HF+SD group (Table 1). Moreover, two of the top upregulated individual genes that we identified were *Adm2*, a vasodilator and angiogenic factor known to respond to TSH stimulation (50–53), and its receptor modulator *Ramp3* (Figure 6A). Immunohistochemistry revealed that overnutrition increased ADM2 expression in thyrocytes (Figure 6, C and D). Thus, we conclude that TSH signaling is increased by overnutrition, not blocked. We also propose that the HF+SD-fed mice exhibit an adaptive mechanism whereby their elevated TSH levels upregulate thyrocyte ADM2 expression, which drives vascularization in a paracrine signaling fashion. Ultimately, this process supports thyroid growth and delivery of nutrients needed for TH biosynthesis, but unfortunately, this compensatory mechanism does not rescue thyroid function.

### Thyroidal changes caused by overnutrition are mostly reversible

An important subsequent question is whether the damage inflicted by overnutrition is reversible. To answer this question, we placed male mice on the HF+SD for 6 weeks and then returned them to the CD for another 6 weeks (the reversed group: REV). The REV mice gained weight and then rapidly lost it, matching the BW, fat mass, and leptin levels of the control mice by the end of the study (Figure 7, A and B; Supplemental Figure 11A), and maintained a normal energy balance during the study’s final weeks (Supplemental Figure 11, B and C), suggesting no modulation of the HPT axis by energy balance during that time. Hearteningly, most of the functional damage to the thyroid induced by overnutrition appears to be reversible by weight loss. Serum T_3_ levels remained unchanged, and T_4_ and TSH levels returned to normal (Figure 7, C–E). The thyroids of the REV mice were only ∼60% larger than those of controls (Figure 7F); this growth more closely resembles the thyroid growth after 3 weeks on the HF+SD than after 6 weeks (∼55% vs. ∼80%; Figure 1H), indicating partial reversal. Whether the thyroids would eventually return to their normal size is unclear. Even so, the thyroidal T_4_ and T_3_ levels and their ratio returned to normal—suggesting that the efficiency of TH biosynthesis was restored—as did the histology (Figure 7, G–I; Supplemental Figure 11, D–G), consistent with the normalization of TSH levels. Furthermore, we found no indication of ER stress, TG and NIS protein levels returned to normal, and the levels of TG’s folding chaperones decreased (Figure 7J; Supplemental Figure 12). These findings strongly suggest that thyroid dysfunction can be reversed by dietary intervention and weight loss in mice.

**Figure 7.**
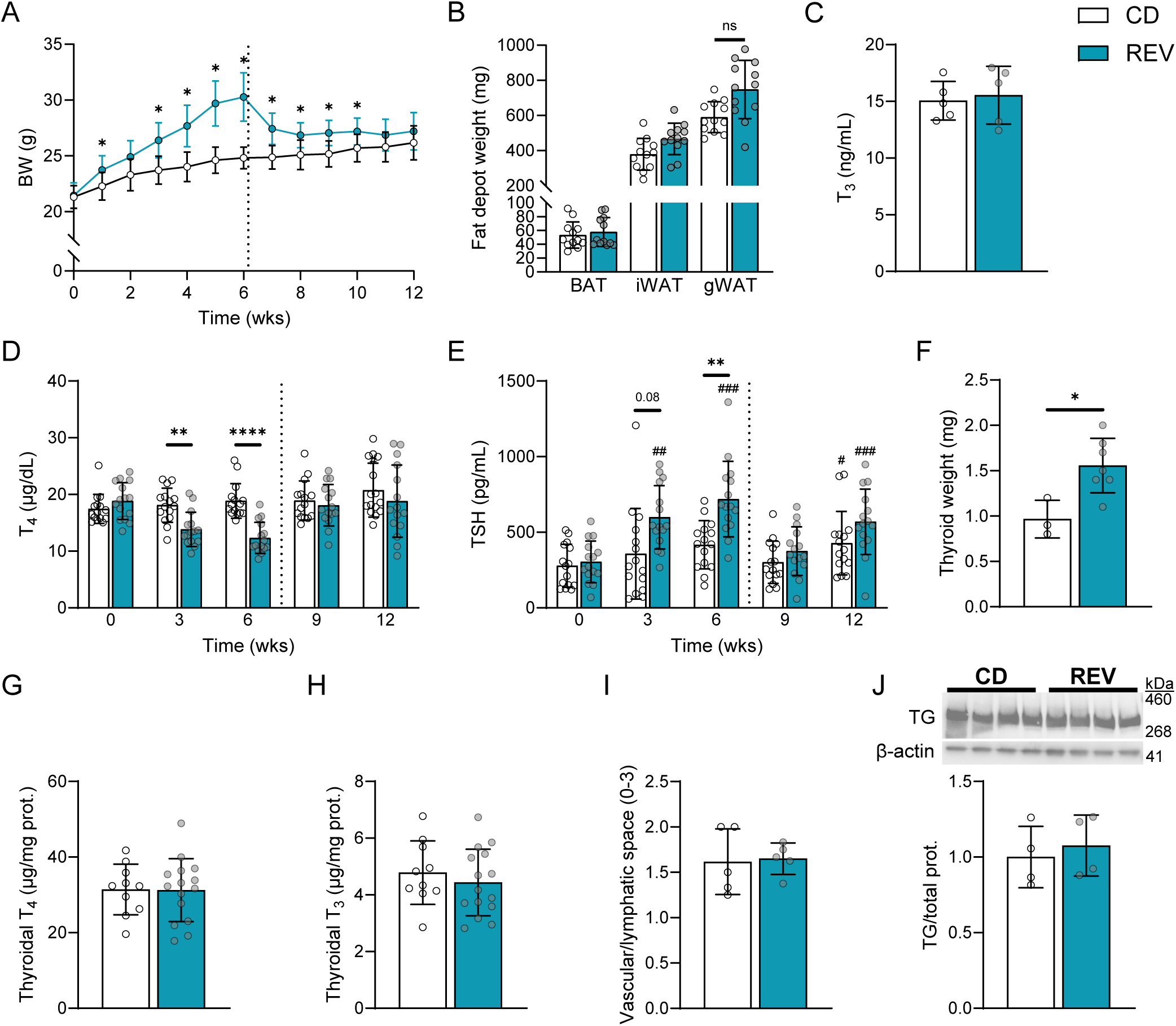
Thyroid dysfunction caused by overnutrition is reversible. Male mice were placed on the HF+SD for 6 weeks and then switched to the CD for 6 weeks (REV). BW was measured weekly (**A**). BAT, iWAT, and gWAT depots were weighed immediately after dissection (**B**). Terminal sera were analyzed by immunoassay for total T_3_ (**C**). Plasma and sera were analyzed by immunoassay for total T_4_ (**D**) and TSH (**E**). Some thyroids were weighed immediately upon dissection (**F**). Other thyroids were fully proteolyzed, and liberated T_4_ (**G**) and T_3_ (**H**) were then measured by immunoassay and normalized to the thyroidal protein. Other thyroids were fixed for H&E staining, and H&E images were then blindly scored for extent of interfollicular vascular/lymphatic space (**I**). Western blot of thyroid homogenate was performed for TG and quantitated (**J**). Data are representative of two separate cohorts of mice: this cohort *n* = 15/group (A), *n* = 12/group (B). Data are results from two separate cohorts of mice per timepoint: combined CD *n* = 10, REV *n* = 15 (G, H). *n* = 15/group (D, E). Dotted line represents switch from HF+SD to CD for REV mice. Data were analyzed by 2-way ANOVA with repeated measures (A, D, E), unpaired Student’s *t*-test with Welch’s correction (B: WAT, C, F–J), or Mann-Whitney test (B: BAT). \**p* < 0.05, \*\**p* < 0.01, & \*\*\*\**p* < 0.0001 vs. controls within that timepoint. #*p* < 0.05, ##*p* < 0.01, & ###*p* < 0.001 vs. week 0 within that diet group.

### Increased BMI is associated with increased thyroidal vascularization in humans

To preliminarily assess the translatability of our findings, we performed histologic analysis of surgically resected human thyroid tissue from patients with multinodular goiter (MNG; Supplemental Figure 13, A–F). We found that patients with obesity (BMI ≥ 30) exhibited increased vascular/lymphatic space, and CD31 staining revealed a stepwise increase in vascularization (Supplemental Figure 13, G–I). Spearman correlation tests between each of these scores and the individual patients’ BMIs revealed that in each case, the score was significantly positively correlated with BMI (Supplemental Table 3). RNA-sequencing results from a subset of these patients showed that *ADM2* did not correlate with BMI, but its closely related family member adrenomedullin (*ADM*) was positively correlated with BMI (Supplemental Figure 14). ADM is also a vasodilator and angiogenic factor and shares a receptor subunit with ADM2 (52). These findings indicate that some of the thyroidal responses to weight gain that we have observed in mice have close counterparts in humans; thus, further investigation is warranted.

## Discussion

The interplay between overnutrition and the HPT axis is extremely complex and varies widely across rodent models. Studies in rats demonstrate activation of the HPT axis mediated by enhanced leptin signaling in TRH neurons and in the melanocortin system, but thyroid status is not uniform (54–58). Moreover, the TRH neurons of C57Bl/6J mice are mostly unresponsive to leptin (59), and additional differences have been documented in the melanocortin system’s response to overnutrition according to rodent strain, diet composition, length of study, and brain region examined (60). Though our findings do not hold for all rodent models, one strength of our mouse model of short-term overnutrition is that C57Bl/6J mice are commonly used in metabolic studies, making our findings broadly relevant. Furthermore, the systemic thyroid status induced—i.e., normal serum T_3_ levels, but decreased serum T_4_ levels, and high TSH levels (Figure 1, E–G)—is characteristic of the early stages of hypothyroidism in humans, when small decreases in serum T_4_ levels stimulate TSH secretion, while other homeostatic mechanisms preserve serum T_3_ levels (61). Since the decreased serum T_4_ levels in our mice are accompanied by increased TSH levels, we conclude that their thyroid status is inadequate. Leptin is not a significant driver of TSH elevation in our model, since leptin levels in the HF+SD-fed mice did not exceed those of controls prior to significant changes in the HPT axis and thyroid histology (Figures 1 and 7; Supplemental Figures 1, 5, A–D, and 11). In line with this, female mice—though resistant to obesity despite increased caloric intake, consistent with well-documented sexual dimorphisms (62)—also exhibited diet-induced hypothyroidism (Supplemental Figure 2), suggesting that their thyroid dysfunction is attributable at least in part to their diet, rather than to increased fat mass, since the female mice do not expand their adipose tissue. More work needs to be done to distinguish the effects of obesity from those of different diets and to determine how they are further affected by sex and length of study.

Even so, we identified remarkable compensatory mechanisms that are induced by overnutrition in both sexes, which are attributable to the elevated TSH levels and would be expected to promote greater TH biosynthesis. Thyrocyte hypertrophy and hyperplasia, i.e., goiter, are classic effects of TSH documented for over a century (2, 43). We are, to our knowledge, the first to report quantitative organelle expansion in thyrocytes during overnutrition (Figure 5; Supplemental Figure 6). Given the requirement for TG and ATP for TH synthesis, it seems unlikely that bigger thyrocytes could synthesize more TH without more ER and mitochondria. Although the underlying mechanism requires further investigation, we believe the organelle expansion we have observed is consistent with the mitogenic effects of TSH. Increased vascularization was also described in goitrous thyroids more than a century ago, in cases of I^-^ deficiency disorder (IDD) (43). Nagasaki et al reported that I^-^-deficient rats exhibited high levels of thyrocyte ADM2 and that TSH stimulates ADM2 secretion in vitro (53). To our knowledge, however, we are the first to report that overnutrition increases vascularization in normal I^-^-sufficient thyroid tissue, accompanied by marked elevation in thyrocyte ADM2 (Figure 6). Thus, while we cannot rule out other possible mechanisms, it is reasonable to conclude that the common cause of increased thyroidal vascularization in IDD and overnutrition is elevated TSH levels.

Despite these thyroidal adaptations triggered by TSH, systemic overnutrition-induced hypothyroidism persists due to direct impairment of thyroid function. We found, for the first time, that overnutrition decreases the T_4_ content of the thyroid within just 10 days (Figure 3). Because thyroid growth does not outpace declines in thyroidal T_4_ content, net T_4_ production from the thyroid gland appears to be reduced by overnutrition. Net T_3_ output from the thyroid may be higher than that of control mice, consistent with the maintenance of serum T_3_ levels despite falling T_4_ levels, whereas thyroidal T_3_ content eventually declines. The T_3_:T_4_ ratio in the thyroid increases dramatically in overfed mice, a clear indication of preferential T_3_ synthesis, an adaptive mechanism for preserving serum T_3_ levels at the expense of the prohormone T_4_ that has been well documented in other contexts involving enhanced TSH stimulation, including IDD (28, 42). TSH promotes preferential T_3_ synthesis by increasing de novo T_3_ synthesis on TG (63) and thyroidal T_4_-to-T_3_ conversion by D1 (64, 65). In our model, TG protein levels are decreased (Figure 5G) even while thyroidal T_3_ levels are maintained, suggesting that perhaps the T_3_ content on TG is increased. We also observed a 4-fold increase in *Dio1* expression in HF+SD thyroids (Table 1). It is likely that these mechanisms are at play in our model of overnutrition-induced hypothyroidism, though we cannot completely exclude other possible mechanisms.

Despite preferential T_3_ synthesis, overall TH biosynthesis is impaired in our mouse model of overnutrition, and we investigated several possible mechanisms to explain this thyrocyte dysfunction. Secretory cells, including thyrocytes, are particularly sensitive to ER stress, because slowed translation during ER stress—though meant to restore proteostasis and prevent apoptosis—can be detrimental to secretory function (66, 67). Indeed, we found that HF+SD-fed males exhibit ER bloating and increased expression of ER stress markers, including phosphorylated eIF2α, alongside reduced TG levels (Figure 5). Few studies have been done on the morphological and biochemical effects of overfeeding on the thyroid, but our results are consistent with those of previous studies carried out in rats (56–58). To our knowledge, just one study of overnutrition’s effects on mouse thyroid function has been previously published (68). All these previous studies attributed diet-induced thyroid dysfunction to lipotoxicity, although only two measured thyroid lipids. Surprisingly, we did not find compelling evidence for ectopic lipid deposition (Supplemental Figure 9). We posit that the rapid production of new membranes to sustain cellular and organellar growth outpaces accumulation of sphingolipids, or that ER stress is limiting ceramide synthesis (49). It is not known whether the small number of LDs we observed would be deleterious for thyrocytes, but lipid deposition has been observed in thyroids resected from some obese patients (68, 69), warranting further investigation. Another possible mechanism for thyrocyte dysfunction is the accumulation of misfolded proteins, including TG, during the prolonged TSH stimulation of thyroid protein synthesis. Many *Tg* mutations result in ER stress and congenital hypothyroidism, some of which resemble the overnutrition-induced ER stress that we and others have observed (67, 70). Additionally, our findings suggest that mitochondrial function may be altered (Supplemental Figures 6 and 7), though information on mitochondrial regulation in thyrocytes is admittedly sparse.

We have also provided evidence that overnutrition decreases intracellular T_4_-to-T_3_ conversion, leading to resistance to T_4_ and reduced EE (Figure 2E). We found no difference in the energetic response to exogenous T_3_ between diet groups, indicating that whole-body T_3_ inactivation rates (either by D3 or D1) are not changed by overnutrition. Because the affinity of D3 and D1 for T_4_ is less than or equal to their affinity for T_3_ (41), it is unlikely that direct inactivation of the exogenous T_4_ by D3 or D1 explains our findings. Thus, we conclude that the effects of overnutrition on the body’s energetic response to THs are mediated by reduced activation of T_4_ rather than by increased deactivation of T_4_—effects likely mediated by reduced peripheral D2 activity in BAT, as suggested by Figure 2C, and potentially in other highly metabolic tissues that express D2, in which D2 is a key regulator of metabolism (1). These findings prompt important questions regarding tissue-specific T_3_ signaling, since both low D2 activity and T_4_ substrate availability could cause local hypothyroidism more severe than the systemic hypothyroidism.

Our finding that decreased D2 activity correlates with decreased EE aligns with the results of previous studies showing that D2KO mice are more susceptible to weight gain, insulin resistance, and cold intolerance (71–73). D2 expression and activity are also inversely correlated with obesity and insulin resistance in humans, in whom D2 is expressed in skeletal muscle, a major contributor to thermogenesis and BMR (74–77). Decreased D2 activity clearly causes metabolic perturbations, but our study showed the converse, pointing to a possible vicious cycle. Because D2 has positive effects on EE, the strategy of promoting endogenous D2 activity to treat obesity has garnered a great deal of interest: this is an attractive approach to enhancing local T_3_ signaling in metabolic tissues while avoiding the harm of systemic hyperthyroidism (78). Several groups have shown that cold exposure recruits BAT activity and increases EE in lean and obese subjects, reducing fat mass in some (79, 80). Other groups have demonstrated that pharmacological induction of D2 with bile acids protects mice from diet-induced obesity and increases EE in humans (81, 82). Conversely, studies showing that weight loss itself can improve BAT activity and that obese subjects exhibit reduced BAT recruitment during cold exposure support the hypothesis that obesity impairs D2 activity (83–85). In line with this, our findings suggest that the dampening of local T_3_ generation by impaired D2 activity, including in BAT, may be an additional pathology related to and exacerbating obesity, lending further plausibility to the notion that D2 activity should be rectified and ideally enhanced to combat obesity. This promising therapeutic approach calls for deeper investigation in the midst of the ever-worsening obesity epidemic, especially given that not all patients respond to GLP-1 receptor agonists (86).

The finding that obesity impairs thyroid function and T_4_-to-T_3_ conversion concurrently is extremely important, since these regulatory processes normally compensate for each other. Primary hypothyroidism is successfully treated with T_4_ monotherapy in most patients, due to the increased production of T_3_ by D2 (87–89). Moreover, mice with defects in deiodinases exhibit normal serum T_3_ levels due to increased TH biosynthesis (90, 91). Our findings in mice, however, are consistent with the hypothesis that diet-induced obesity delivers a double blow to TH action by impairing both thyroid function and peripheral D2 activity, likely limiting compensation. The impairment of T_4_ utilization by obesity provides a reasonable explanation for why treating obese subjects with T_4_ does not consistently cause them to lose weight (92). Only supraphysiological doses of TH cause weight loss, but they cannot be administered safely (78). T_3_ mimetics, however, bypass D2, and T_3_ therapies targeted to the liver have been successful (93), most notably a recently FDA-approved TH receptor β agonist effective against MASLD (94). Additionally, T_3_ delivery targeted to adipose tissue decreased BW and improved metabolic function in several mouse models of diet-induced obesity without major side effects (95). Clearly, enhancing T_3_ signaling is a promising approach that should be pursued further, particularly because peripheral T_4_-to-T_3_ conversion decreases during calorie deficit to conserve energy in lean and obese subjects (96–98), explaining, in part, why dieting is largely ineffective (39, 99). Moreover, there is evidence that deiodinase dysregulation persists after weight loss (100) and may contribute to patients’ well-documented propensity to regain weight. Thus, subjects striving to maintain weight loss may still require thyroid support. Likewise, though weight loss itself completely rescues the function of the thyroid gland in mice (Figure 7; Supplemental Figures 11 and 12), inspiring hope for similar outcomes in humans, longer-term metabolic disease may cause lasting thyroid damage.

Importantly, then, we have shown—albeit very preliminarily—that high BMI is correlated with thyroidal vascularization in patients with MNG (Supplemental Figure 13; Supplemental Table 3), similarly to what occurs in our mouse model of diet-induced obesity. *ADM*, which is closely related to *ADM2*, may be a driving factor in these patients, further suggesting that our findings may be translatable (Supplemental Figure 14). Well-preserved normal thyroid tissue is scarce, but this study should be repeated in healthy subjects whose TSH levels can be ascertained. A second limitation of the gene expression study is that the sample size was small (*n* = 20) and consisted almost exclusively of obese subjects. Thus, it is possible that *ADM2* will be correlated with BMI when subjects of normal weight are included. In fact, *ADM2* was recently reported to be associated with high BMI and tumor aggression in patients with thyroid cancer (101). More work is needed to understand these associations and to determine whether biochemical features of overnutrition-induced thyroid impairment in mice also occur in humans with high BMI who are otherwise healthy.

The work presented here begins to fill important gaps in our knowledge using a frequently employed mouse model of diet-induced obesity. This model recapitulates the thyroid phenotype present in obese patients: increased thyroid volume, TSH levels, and serum T_3_:T_4_ ratio, symptoms routinely interpreted as indicating thyroid dysfunction in other contexts (19, 22, 23, 25, 26). Our findings are in line with previous research demonstrating that thyroid dysfunction can be induced by overnutrition, and adds important elements such as quantitative EM analysis, direct measurement of thyroidal THs, and metabolic analysis of TH action. Some of these changes appear to be driven by elevated TSH levels, or possibly by ectopic lipid deposition, or by both; these are phenomena that occur in a subset of obese subjects (22, 68, 69). Taken together, our findings support the emerging hypothesis that some patients may have hypothyroidism secondary to obesity, and call for deeper investigation of obesity-induced thyroid dysfunction in humans. Moreover, a recent bidirectional Mendelian randomization analysis revealed that genetically predicted high BMI was significantly associated with increased TSH levels, but not conversely (102). This issue is highly important, given the metabolic dysfunctions associated with hypothyroidism, including reduced EE (6–10), which can exacerbate obesity. Crucially, hypothyroidism occurs early during overnutrition in our model, with signs of thyroidal stress apparent after only 10 days of overnutrition and worsening over time. This outcome has important ramifications for the many other studies using mouse models of diet-induced obesity that often last 20 weeks or longer, in which, surprisingly, THs are rarely considered. Our work furnishes a strong argument for including TH analysis in these studies, particularly for tissues that rely on serum T_4_ to generate intracellular T_3_. Fortunately, thyroid dysfunction in mice can be reversed by weight loss, offering hope that the same strategy could be used to repair obesity-induced thyroid damage in patients, thereby improving their metabolic regulation. Our study may help establish the basic thyroidal pathophysiology that we must understand in order to guide pharmacological interventions for improving thyroid function in obese patients, which will need to involve clever approaches to improving or bypassing deiodinase function.

## Materials and Methods

### Sex as a biological variable

Preliminary mouse studies were performed in both sexes. The overall phenotype (hypothyroidism, goiter, thyroid histology, etc.) was similar between sexes, so later studies were performed with only male mice. Studies with human subjects incorporated both sexes; BMI categories were sex-matched. Data were analyzed together because male subjects were few, due to the much higher prevalence of thyroid disease among females (2).

### Animals

C57Bl/6J mice were purchased (Jackson) and then bred in-house. Mice were co-housed unless otherwise noted. When 6-7 weeks old, mice were started on the study chow diet (CD; Inotiv #TD.190244) to acclimate. At 8 weeks of age, about half the mice were placed on high-fat diet (HFD; 60% kcal from lard; Inotiv #TD.190243) and 5% sucrose water (collectively, the HF+SD). Sucrose water was prepared every 1-2 weeks, sterile filtered, and kept at 4°C. The HFD and CD are micronutrient-matched. For the reversibility studies, the mice on the HF+SD were switched back to CD after 6 weeks. BW and each cage’s food and water consumption were measured weekly. All mice were housed at 23°C on a 12-h light cycle.

### Human subjects

Surgically-resected FFPE thyroid tissues were obtained from de-identified patients treated for MNG at Vanderbilt University Medical Center (VUMC).

### Tissue & blood collection

At the end of each study, mice were euthanized by exsanguination with removal of a vital organ under deep isoflurane anesthesia. Organs and tissues were collected, fat depots and thyroids were weighed, and all tissues were snap frozen. Mice from each experimental group were rotated to randomize the time of tissue collection. Tissues were stored at -80°C until analysis.

Survival bleeds were performed by the retro-orbital (RO) method under deep isoflurane anesthesia using heparin-coated capillary tubes. RO blood collection was performed 9AM-12PM to minimize possible circadian effects. End-point bleeds were performed by RO, or blood was collected via syringe during exsanguination. Blood was allowed to sit at room temperature (RT) for at least 30 min before spinning down (5000 rpm, 10 min, 4°C). Plasma and sera were then transferred to a fresh microcentrifuge tube and stored at -80°C until analysis.

### TH & TSH quantitation

For plasma and sera, samples were diluted 1:10 in 1X PBS. T_4_ and T_3_ were measured by immunoassay using commercial kits (Diagnostic Automation/Cortez Diagnostics, Inc. #9003-16 & #9001-16, respectively). TSH was measured neat by immunoassay using a commercial kit (Millipore #MPTMAG-49K). All were done according to the manufacturers’ instructions.

For thyroids, proteolytic digestion was performed to degrade TG and liberate the THs for quantitation, following a slightly modified published protocol (103). Briefly, deep-isoflurane-anesthetized mice were perfused transcardially with 35°C 0.9% saline for 1 min to remove blood-borne THs. Thyroids were excised quickly, flash frozen in liquid nitrogen, and stored at -80°C until analysis. Next, one lobe was power homogenized in homogenizing buffer (100 mM Tris-HCl, pH 8.6; 50 mM sodium azide; 50 mM EDTA). Samples were spun down, and 150 µl supernatant was transferred to a tube containing digestion buffer and toluene. Remaining supernatant was stored for protein quantification by the BCA method to normalize results. Digested sample T_4_ and T_3_ were measured with commercial kits (described above). Samples were diluted ∼100-fold for T_4_ and ∼800-fold for T_3_ in digestion buffer lacking Pronase (10 parts) and toluene (1 part) so that readings fell within the linear part of the standard curve.

### Deiodinase activity assays

The deiodinase activity assays were performed in compliance with American Thyroid Association (ATA) guidelines (104). Tissues were homogenized in PE-sucrose buffer (1 mM EDTA, 0.25 M sucrose, 1X PBS without Ca^2+^/Mg^2+^) on ice. Sample protein concentrations were determined by the BCA method; DTT (to 20 mM) was then added to each sample. D1 activity in liver homogenates (250 µg protein) was assayed at 37°C with T_4_ reaction solution (0.5 nM cold T_4_, 20 mM DTT, 115,000 cpm [^125^I]T_4_, PE-sucrose buffer). BAT D2 activity was assayed under the same conditions but adding 1 mM PTU and 10 nM cold T_3_. After 0.5-4 h, reactions were stopped with horse serum. Protein and iodothyronines were precipitated with 50% TCA and spun down. Free ^125^I^-^ in the supernatants was then measured with a gamma counter. D1 and D2 activities were calculated as I^-^ produced/protein content/time. D3 activity in hippocampus and cortex homogenates (25 µg protein) was assayed at 37°C with T_3_ reaction solution (1 nM cold T_3_, 20 mM DTT, 160,000 cpm [^125^I]T_3_, PE-sucrose buffer). Reactions were stopped with methanol and spun down to pellet proteins. Supernatants were then filtered and passed through an ultra-HPLC with an in-tandem flow scintillator detector to separate and count [^125^I]T_2_ and [^125^I]T_3_. D3 activity was calculated as [^125^I]T_2_ produced/protein content/time. Prior to each assay, the outer ring-labeled [^125^I]TH substrates were purified by Sephadex column.

### Metabolic cage analysis

Male mice were kept on either HF+SD or CD for 4 weeks and then placed in individual metabolic cages (Sable Promethion Core) for 1 week (Figure 2D). Mice were acclimatized to single housing the week prior and continued on their respective diets throughout. The first 48 h of metabolic cage data (acclimatization) were discarded. Data were subsequently collected for 48 h to determine each mouse’s baseline EE. Mice were then implanted with an osmotic minipump (Alzet #2001) containing either T_3_, T_4_, or vehicle. T_3_ and T_4_ stock solutions (1.4 mM and 8 mM, respectively) were prepared in 40 mM NaOH per ATA guidelines (104) and sterile filtered. Stock solutions were diluted in sterile 0.9% saline such that final delivery rate was 1.5 µg T_4_/g BW or 0.11 µg T_3_/g BW per day. Minipumps were loaded according to the manufacturer’s instructions and implanted subcutaneously above the interscapular BAT under deep isoflurane anesthesia. Mice were administered analgesia (10 mg/kg BW ketoprofen) and then returned to the metabolic cages. After one recovery day, each mouse’s EE was measured over two 24-h time periods (“Day 1” & “Day 2”) and compared to its baseline to calculate percent change. The averages for the baseline, Day 1, and Day 2 periods used to calculate percent change were determined using CalR (105). EE was calculated using the Weir equation. Because data for each mouse were normalized to that mouse’s own baseline, data normalization to BW or composition was unnecessary.

### Fixed tissue collection for histology

Deep-isoflurane-anesthetized mice were perfused transcardially with ice-cold 1X PBS for 5-10 min to wash out blood and then fixed with 4% paraformaldehyde in 1X PBS for 5-10 min. Fixed tissues were dissected and stored in 4% paraformaldehyde at 4°C at least overnight, transferred to 70% ethanol, and then stored at RT until paraffin embedding and sectioning (5 µm).

### Histological staining

Fixed mouse thyroid sections were stained with either H&E or α-mouse CD31 antibody (Dianova #DIA310) and hematoxylin using an automated stainer (Sakura Tissue-Tek Prisma Plus and Leica Bond-RX, respectively). Some H&E stains were performed manually following the standard protocol. Slides were imaged with a scanning brightfield microscope (Leica). De-identified screenshots of H&E stains (2-5 fields/animal in most cases) and CD31 stains (1 field/animal) were blindly scored by two independent thyroid pathologists.

ADM2 staining of fixed mouse thyroid sections was performed manually. Briefly, slides were de-paraffinized and rehydrated. Antigen retrieval was then performed in boiling citrates buffer (10 mM, pH 6.0). Slides were pre-treated with 3% hydrogen peroxide, blocked [4% BSA (w/v), 0.3% Triton X-100, 1X PBS], and then incubated at 4°C overnight with α-ADM2 antibody (Invitrogen #PA5-72030) in blocking solution. Slides were then incubated with HRP-conjugated secondary antibodies, developed with DAB substrate solution (Vector Labs #SK-4100), and counterstained with hematoxylin. Finally, slides were dehydrated, and coverslips were mounted with resin. Negative controls were treated the same way but without primary antibody. Slides were imaged with a brightfield microscope (Leica).

### Scanning electron microscopy (SEM)

Deep-isoflurane-anesthetized mice were perfused transcardially with 35°C 1X PBS without Ca^2+^/Mg^2+^ for 30 sec at 8 mL/min to wash out blood and then fixed with 35°C SEM fixative buffer (2% glutaraldehyde, 2% paraformaldehyde, 2 mM CaCl_2_, 0.1 M sodium cacodylate) for 5 min (106). The thyroid was then dissected, cut into ∼1 mm^3^ pieces, and postfixed in a second SEM fixative buffer (2.5% glutaraldehyde, 2 mM CaCl_2_, 0.1 M sodium cacodylate) for 1 h at RT. Thyroids were transferred to 1X PBS and stored at 4°C overnight. Further processing was performed as described (107). SEM-ready thyroids were cut into 100 nm-thick sections using an ultramicrotome (Leica) and placed on 5x7 mm silicon wafers (Electron Microscopy Sciences) prior to SEM imaging. Thyroid follicles were identified by their classical anatomy of a colloid-filled lumen surrounded by multi-ciliated epithelial cells (i.e., thyrocytes). Thyrocytes were imaged by a Crossbeam 550 (Zeiss) operating at 3 keV and 1 nA, guided by automated tile acquisition and stitching using the Atlas 5 software (Zeiss). SEM images have an X-Y resolution of 5 nm, and at least 6 follicles/animal were imaged.

### SEM image analysis using machine learning

To quantify thyrocyte organelle composition and anatomic parameters, we created 2D U-nets trained to identify and segment ER and mitochondria compartments. Individual thyrocytes with a visible nucleus and apical contact with colloid were manually segmented. U-nets were trained using Aivia software (Leica Microsystems). ER or mitochondria U-nets were trained using up to 30 individual and representative areas of thyrocyte cytosol that contained ER and mitochondria compartments. Individual ER and mitochondria were manually annotated using the LabKit plugin (ImageJ) and loaded into the U-net. The following training parameters were used: 8 layers, 64 Init Filters, 64 Filter Growth Factor, a channel reduction factor of 8, an image block size of 256x256 pixels, and an intensity threshold and area ratio threshold of 0.05. The Adam optimizer with a learning rate of 0.0001 and a staircase exponential decay for the learning rate scheduling method was used. The ER model was trained using 10,000 epochs, and the mitochondria model was trained using 600 epochs. In both models, each epoch contained 256 steps, and the balanced binary cross entropy loss function was used. The resulting trained models were applied in batch to SEM images of single thyrocytes manually segmented using ImageJ. Trained models created 32-bit pixel “organelle probability maps” that were thresholded to include only pixels with ≥80% or ≥95% classification confidence for ER and mitochondria, respectively. Nucleus area was manually annotated (LabKit) and subtracted from each image; additional post-hoc processing was conducted to improve model performance. Processed images were used to quantify relative organelle composition in each cell. Nucleus area was subtracted from total cell area to calculate cytosol area. CellProfiler’s “*MeasureAreaOccupied*” function was used to determine the relative fraction of cytosol occupied by ER or mitochondria objects.

### Western blotting

Frozen thyroid tissue was homogenized in either 1X PBS supplemented with protease and phosphatase inhibitors (Pierce #A32959 or Halt #87786) in lysis buffer (50 mM Tris-HCl, pH 7.5; 150 mM NaCl; 5 mM EDTA; 1% Triton X-100) or in RIPA lysis buffer (Sigma #R0278) supplemented with protease and phosphatase inhibitors. Samples were spun down, and supernatants were taken for further analysis. Samples for NIS blots were heated at 37°C for 30 min, not boiled as the other samples. Proteins (5-20 µg) were separated by SDS/PAGE (Bio-Rad, Invitrogen), transferred to nitrocellulose membrane, blocked (5% BSA, 0.1% Tween-20, 1X TBS), and probed with primary antibodies (Supplemental Table 4). Some blots were stripped with stripping buffer (homemade: 0.2 M glycine, 1% SDS, 0.1% Tween 20, HCl to pH 2.3; or commercial: Thermo #TS46436) and reprobed. Signal was developed by Pico PLUS or Femto Maximum (Thermo #34580 and #34094, respectively) ECL substrate, imaged with a ChemiDoc imaging system (Bio-Rad), and quantified using Image Lab software (Licor). Total protein imaging was done with either Ponceau S stain or stain-free technology (Bio-Rad) prior to blocking. Because TG is 50% of the protein in the thyroid (2) and it was changed by overnutrition, proteins of interest were normalized to total protein below the TG fragment bands (below ∼150 kDa).

### RNA sequencing

RNA was extracted (Trizol/Qiagen RNeasy) from flash-frozen mouse thyroids, cDNA libraries were constructed using the NEBNext Ultra II RNA Library Prep Kit (NEB #E7765L), and next-generation sequencing (Illumina NovaSeq 6000) was performed, according to the manufacturers’ instructions. Data were processed using the DESeq2 package. Detailed protocols are available in the GEO database (accession #GSE294396).

### Statistical analysis and figures

Results were analyzed with GraphPad Prism software as described individually in the figure legends. Normality of data sets was tested to aid selection of the appropriate parametric or non-parametric statistical test. Data are presented as mean ± SD. Schematic figures were created with BioRender.

### Study approval

All mouse experiments were carried out in accordance with VUMC’s IACUC. All studies with human subjects were approved by VUMC’s IRB. This is a retrospective cohort, and it is not possible to consent these patients with historic samples.

### Data availability

All data from mouse studies are available in the Supporting Data Values file, except mouse RNA-sequencing data, which is available in the GEO database (accession #GSE294396). There are restrictions to the availability of patient clinical and sequencing data. Because it is not possible to retrospectively consent these patients, the IRB has requested that we not publicly share individual-level data. The data are securely stored within a Vanderbilt patient data system. Aggregate-level data reported in this paper will be shared by Dr. Weiss upon request. Individual-level data are available only through collaboration following approval of Dr. Weiss and VUMC’s IRB.

## Author Contributions

JR and NC conceived the project. QS and MAL performed the human RNA-sequencing analysis; MAL performed the corresponding Pearson correlations. FSL and ACB carried out and advised on the deiodinase activity assays. RAD developed the deep learning models; JR performed the associated manual segmentation. HW and VLW performed the histological scoring. VLW provided and oversaw access to human data. JR performed all other experimental work, with assistance from APTM and KH. JR, APTM, and NC wrote the manuscript with input from all other authors.

## Acknowledgements

We thank the members of the Carrasco laboratory for critical reading of the manuscript and insightful discussion. We thank Dr. Daniel Canals and Dr. Yusuf Hannun (Stony Brook University) for performing the lipidomics. We also thank the following core laboratories at Vanderbilt University or VUMC for their excellent support: the Translational Pathology Shared Resource (NCI/NIH Cancer Center Support grant P30CA068485; Shared Instrumentation grant S10 OD023475-01A1), the Digital Histology Shared Resource, the Cell Imaging Shared Resource (NIH grants CA68485, DK20593, DK58404, DK59637, and EY08126; Shared Instrumentation grant S10OD028704-01A1), the Mouse Metabolic Phenotyping Center (NIH grants DK135073 and DK020593), the Analytical Services Core (NIH grant DK020593), Vanderbilt Technologies for Advanced Genomics, Creative Data Solutions, and the Molecular Cell Biology Resource. This study was supported by Vanderbilt Diabetes Research & Training Center Pilot and Feasibility grant DK020593 (NC); Vanderbilt Institute for Clinical and Translational Research grant VR70041 (JR); NIH grants VCORCDP K12CA090625, K08CA240901, and R01CA272875 (VLW); NIH grants F30CA281125-01 and T32GM007347 (MAL); American Cancer Society grants RSG-22-084-01-MM and 133934-CSDG-19-216-01-TBG (VLW); and ATA grant 2019-0000000090 (VLW).

The NIH Public Access Policy grants the NIH the right to make this federally funded work publicly available in PubMed Central.

## Notes

### Competing Interest Statement

Only Dr. Bianco has disclosures to declare, namely, that he has consulted for AbbVie, Acella, Aligos, and Synthonics. His work as a consultant had no bearing on the results presented here. The other authors have declared that no conflict of interest exists.

### Summary of Updates

Feedback from journal peer reviewers has been incorporated, including two panels added to the supplemental material (Supplemental Figures 4D and 11A), movement of all studies with human samples to the supplemental material (now Supplemental Figure 13), and minor text revisions throughout.

