## Supplemental Material for "Overnutrition directly impairs thyroid hormone biosynthesis and utilization, causing hypothyroidism, despite remarkable thyroidal adaptations"

**Supplemental Materials and Methods:**

### *Urinary $I^-$ analysis*

Mice were singly housed in urine collection cages for 48 h with *ad libitum* access to food and water. A layer of mineral oil was added to the collection tubes to block evaporation. Every 24 h, urine was aspirated from under the mineral oil and spun down to remove particulates (3000 x g, 10 min, 4°C). Supernatants were stored at -20°C until analysis. $I^-$ concentration was measured by colorimetric assay (BioVision #K2037-100) according to the manufacturer's instructions.

### *Leptin measurement*

Blood was collected and processed as described in the main text. Plasma and serum samples were diluted 1:5 in diluent (0.05% Tween-20; 0.1% BSA; 1X PBS, pH 7.2; sterile filtered). Leptin was measured by ELISA using a commercial kit (PeproTech #900-K76) according to the manufacturers' instructions.

### *Western blotting*

Frozen BAT was homogenized in lysis buffer (50 mM Tris-HCl, pH 7.5; 150 mM NaCl; 0.5% IGEPAL; 1 mM $Na_4P_2O_7$; 1 mM benzamidine; 5 mM $Na_3VO_4$; 10 mM NaF) supplemented with protease inhibitors (Sigma Roche #04693159001). Samples were spun down, and the non-fatty phase was taken for further analysis. Thyroids were processed as described in the main text. Samples for OxPhos complexes and NIS blots were heated at 37°C for 30 min or 10 min, respectively, not boiled as the other samples. Proteins (5-20 µg) were separated by SDS/PAGE (Bio-Rad), transferred to nitrocellulose or PVDF membrane, blocked (5% BSA, 0.1% Tween-20, 1X TBS), and probed with primary antibodies (Supplemental Table 3). Some blots were stripped with commercial stripping buffer (Thermo #TS46436) and reprobed. The signal was developed by Pico PLUS (Thermo #34580) or Femto Maximum (Thermo #34094) ECL substrate, imaged with a ChemiDoc imaging system (Bio-Rad), and quantified using Image Lab software (Licor). Total protein imaging was done with stain-free technology (Bio-Rad) prior to blocking. Because TG is 50% of the protein in the thyroid (2) and it was changed by overnutrition, proteins of interest were normalized to total protein below the TG fragment bands (below ~150 kDa).

### *Triglyceride quantitation*

Frozen thyroid tissue was homogenized in 1X PBS. Triglycerides were then measured using a commercially available bioluminescent assay kit (Promega #J3160) according to the manufacturer's instructions.

### *Lipidomics*

Lipid extraction and analysis by HPLC-electrospray tandem mass spectrometry of frozen thyroid tissue were carried out according to previously published protocols (3). Inorganic phosphate was quantitated to normalize results.

### *RNA-sequencing analysis*

Mouse RNA-sequencing data were collected as described in the main text. GSEAs were performed in R with MSigDB gene set collections.

Human RNA-sequencing data were derived from FFPE thyroid tissue samples from patients treated for MNG at VUMC. RNA extraction, sequencing, and analysis of tissue samples from this cohort was previously described (4), from which DESeq2 normalized gene counts were used.

### *Histological staining*

H&E-stained slides of FFPE human thyroid tissues were retrieved from VUMC storage. Sections from the associated blocks were cut and stained with α-human CD31 antibody (Leica #PA0250) and hematoxylin using an automated stainer (Leica Bond-RX). Slides were examined under a microscope and blindly scored by two independent thyroid pathologists. Representative images were taken.

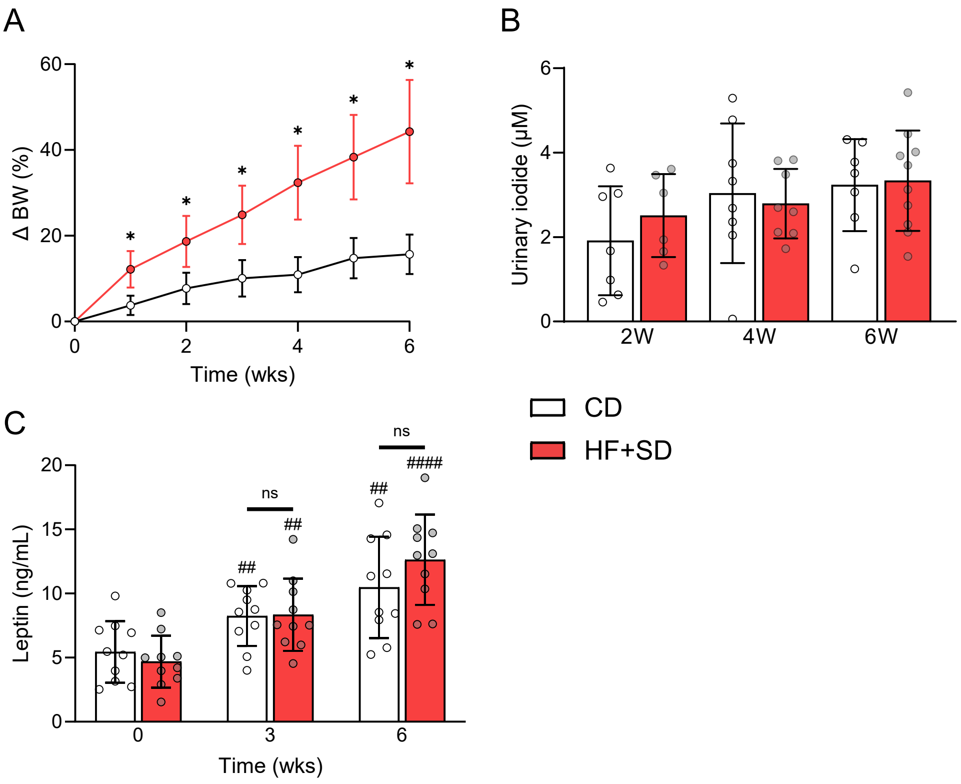


**Supplemental Figure 1. Weight gain-associated changes in the HPT axis are not due to I⁻ deficiency or increased leptin.** Male mice were placed on the HF+SD or CD for 6 weeks, and their cumulative percent BW change from baseline was calculated per mouse (**A**). Urine was collected and its $I^-$ concentration was measured by colorimetric assay (**B**). Plasma and sera were analyzed by immunoassay for leptin (**C**). Data are representative of at least two separate cohorts of mice: this cohort CD $n$ = 20, HF+SD $n$ = 25 (A). Data were analyzed by 2-way ANOVA with repeated measures (A, C) or at each individual timepoint by unpaired Student's *t*-test with Welch's correction (B) since each timepoint is a separate cohort of mice. *$p$ < 0.05 vs. controls within that timepoint. ##$p$ < 0.01 & ####$p$ < 0.0001 vs. week 0 within that diet group.

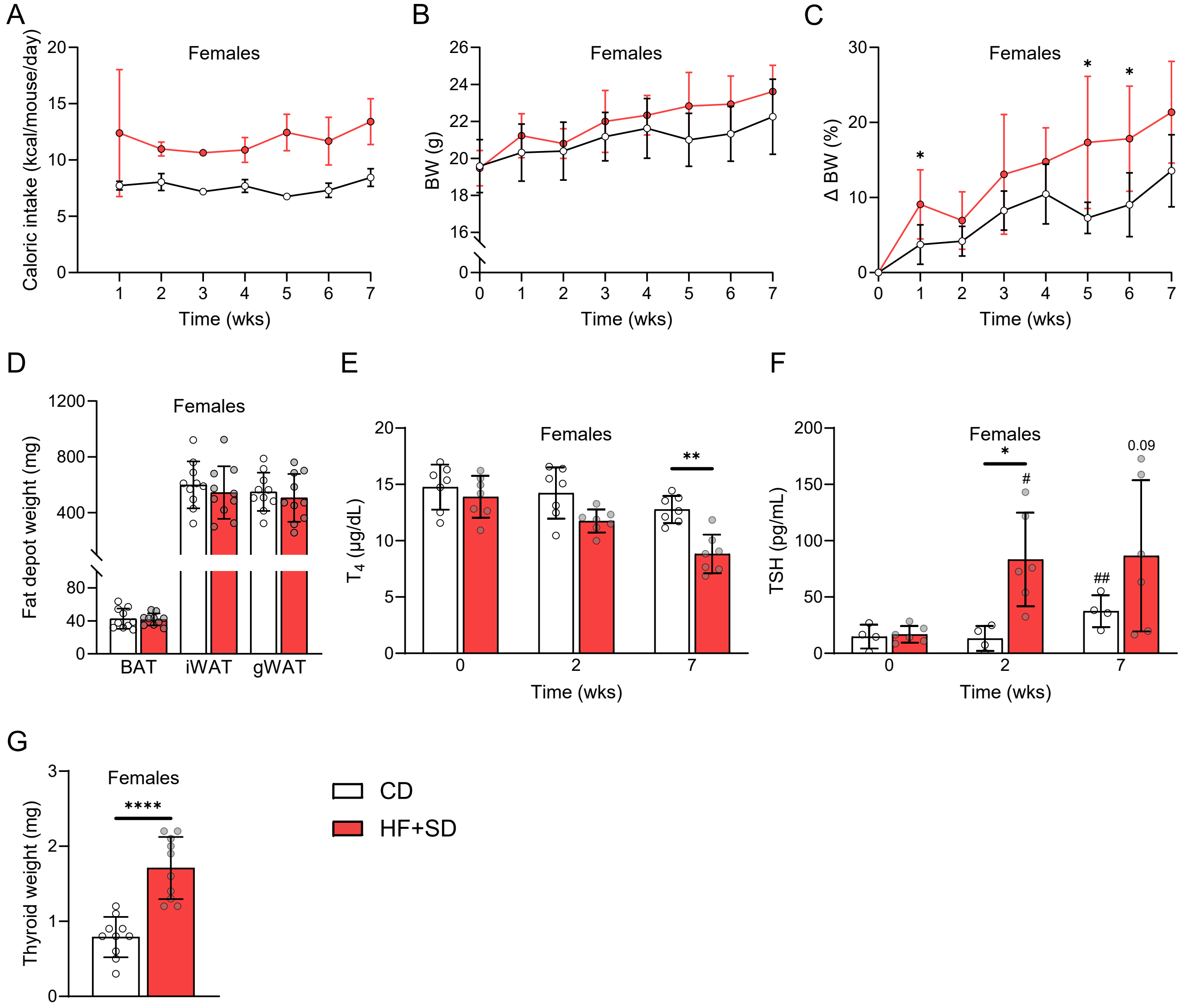


**Supplemental Figure 2. Overnutrition induces mild hypothyroidism in female mice.** Female mice were placed on the HF+SD or CD for 7 weeks and their caloric intake (**A**) and BW (**B**) were measured weekly. Cumulative percent BW change from baseline was calculated per mouse (**C**). BAT, iWAT, and gWAT depots were weighed immediately after dissection (**D**). Plasma and sera were analyzed by immunoassay for total $T_4$ (**E**) and TSH (**F**). Thyroids were carefully dissected and weighed (**G**). $n$ = 10/group in 2 cages/group (A–D). Caloric intake data were not statistically analyzed because $n < 3$/group (A). Other data were analyzed by 2-way ANOVA with repeated measures (B, C, E, F) or unpaired Student's *t*-test with Welch's correction (D, G). *$p < 0.05$, **$p < 0.01$, & ****$p < 0.0001$ vs. controls within that timepoint. #$p < 0.05$ & ##$p < 0.01$ vs. week 0 within that diet group.

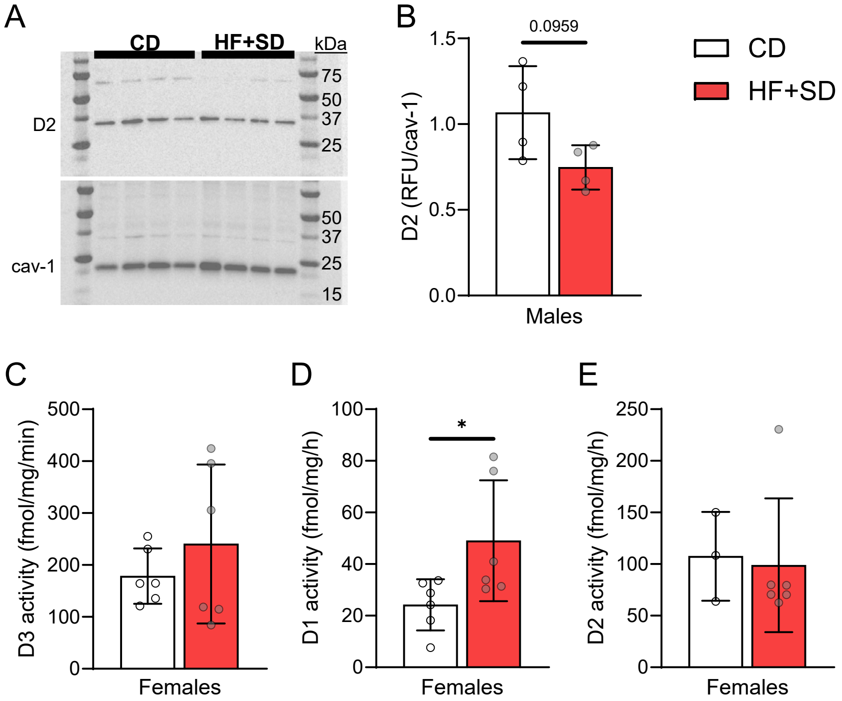


**Supplemental Figure 3. Overnutrition modulates the deiodinases similarly between sexes.** Male mice were placed on the HF+SD or CD for 6 weeks. BAT homogenates were probed for D2 by Western blot and reprobed for the housekeeping protein caveolin-1 (**A**), and the results were quantitated (**B**). Female mice were placed on the HF+SD or CD for 7 weeks. Deiodination rates mediated by D3 in cortex (**C**), D1 in liver (**D**), and D2 in BAT (**E**) were measured. Data were analyzed by unpaired Student's *t*-test with Welch's correction. *$p < 0.05$.

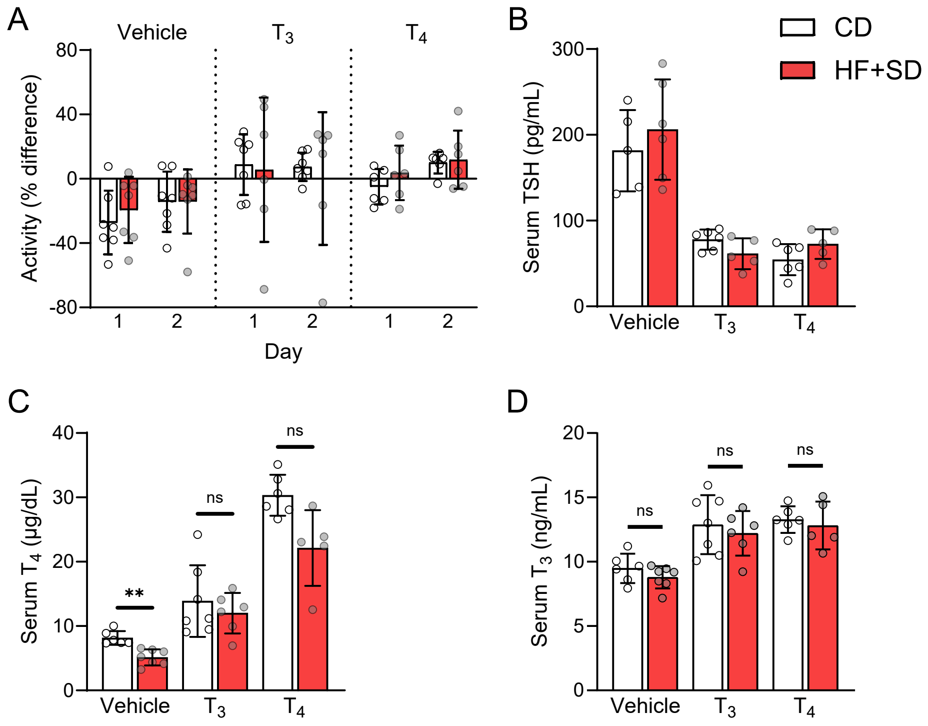


**Supplemental Figure 4. Differences in EE were not due to differences in serum TH levels or physical activity levels.** Male mice were placed on the HF+SD or CD for 4 weeks, and then metabolic cage analysis was performed before and after implantation of osmotic minipumps delivering THs or vehicle. Activity was measured at baseline for 48 h and in response to vehicle, $T_3$, or $T_4$ for two 24-h time periods ("Day 1" and "Day 2") after osmotic minipump implantation. The percent difference of the average activity between the response "Days" and baseline was then calculated for each mouse (**A**). Serum TSH (**B**), total $T_4$ (**C**), and total $T_3$ (**D**) were measured by immunoassay at the end of the study. Data were analyzed by 2-way ANOVA with repeated measures (A) and Brown-Forsythe and Welch ANOVA tests (B–D). $^{**}p < 0.01$.

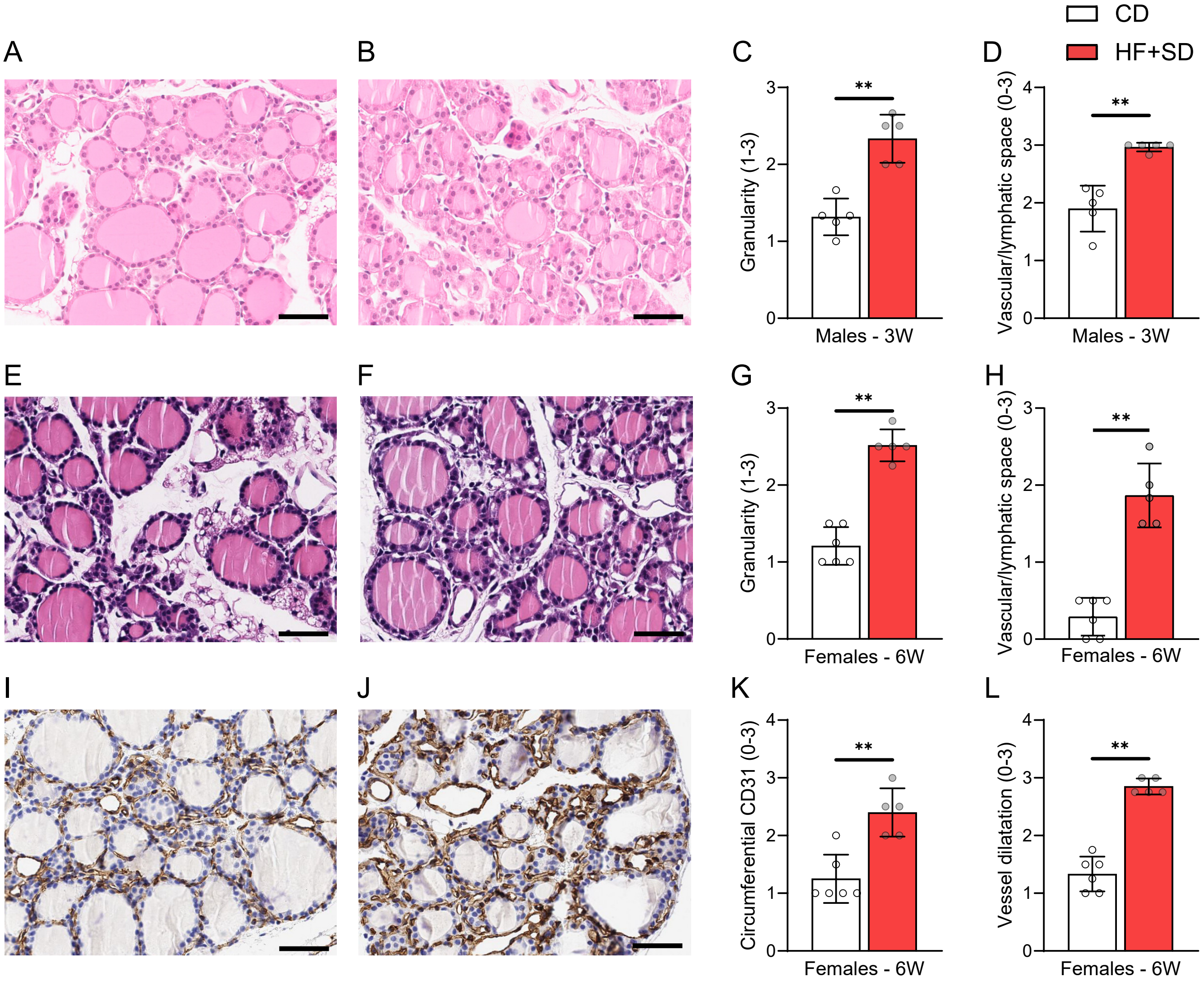


**Supplemental Figure 5. Overnutrition increases thyrocyte size, granularity, and vascularization in female mice and within just 3 weeks in male mice.** Male mice were placed on the HF+SD or CD for 3 weeks. Representative H&E images from a CD mouse (**A**) and a HF+SD mouse (**B**) are shown. H&E images were blindly scored for extent of granularity (**C**) and interfollicular vascular/lymphatic space (**D**). Female mice were placed on the HF+SD or CD for 6 weeks. Representative H&E images from a CD mouse (**E**) and a HF+SD mouse (**F**) are shown. H&E images were blindly scored for extent of granularity (**G**) and interfollicular vascular/lymphatic space (**H**). Their fixed thyroid sections were also stained for CD31, a vascular marker. Images from CD mice (**I**) and HF+SD mice (**J**) were blindly scored for extent of circumferential staining around the follicles (**K**) and dilatation of the follicular microcapillaries (**L**). Scale bars = 50 μm. Data were analyzed by Mann-Whitney test. ***p* < 0.01.

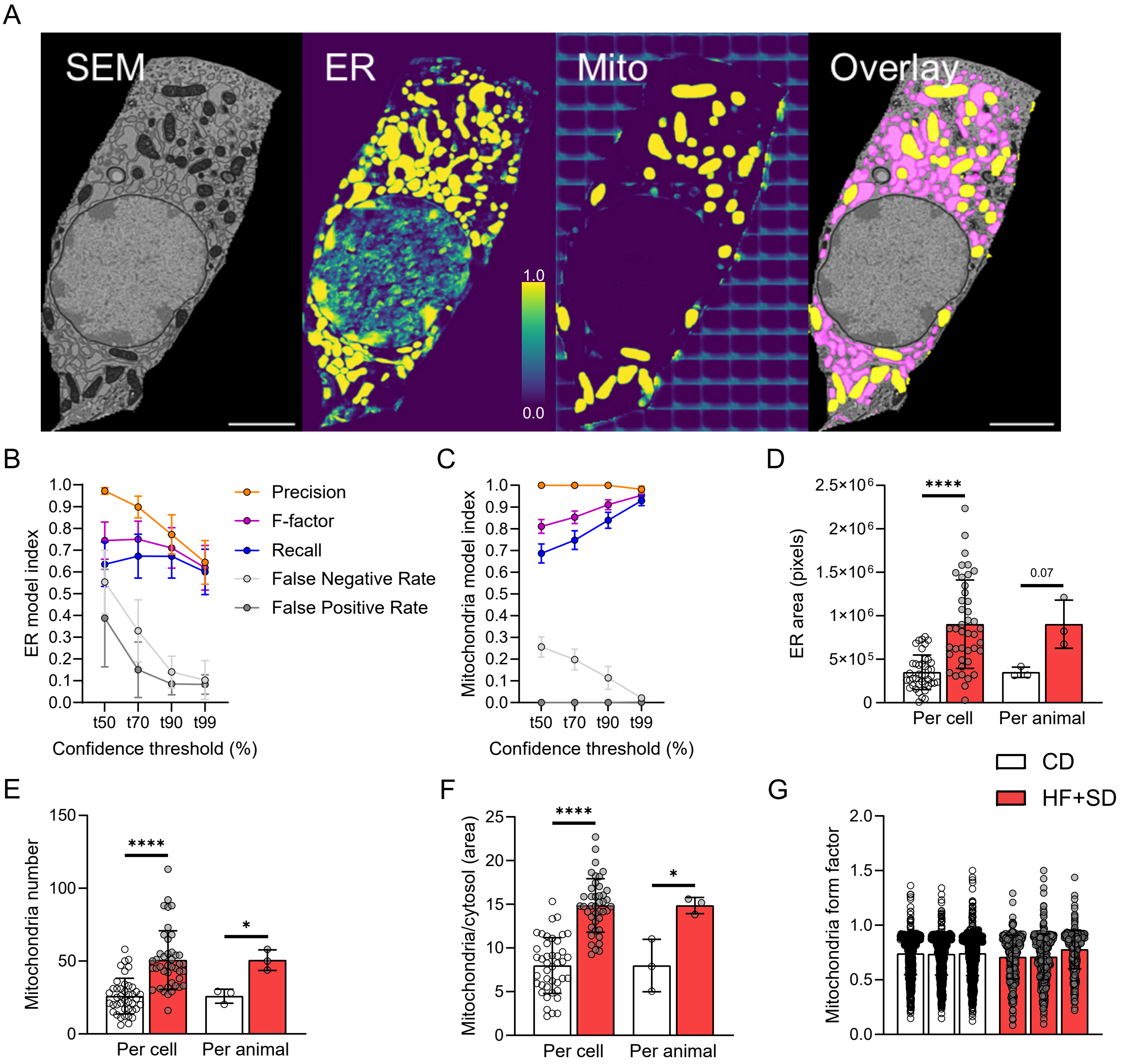


**Supplemental Figure 6. The deep learning models achieved a high degree of accuracy and revealed that overnutrition expands ER area and mitochondrial density.** Male mice were placed on the HF+SD or CD for 6 weeks. Thyroids were processed for EM. Deep learning models were trained to segment the ER and mitochondria (**A**). Pixel-level evaluation indices based on the training set images are shown for the ER (**B**) and mitochondria (**C**) models. A t value of 80% and 95% was chosen for ER and mitochondria, respectively. Post-hoc processing, including subtraction of manually segmented nuclei from each image, was performed to improve the accuracy of the models, which were then used to quantitate ER area (**D**), the number of mitochondria (**E**), and mitochondrial area relative to cytosol area (**F**). The degree of mitochondrial branching (form factor) was also calculated (**G**). Scale bars = 2.5 μm. Model index data are presented as mean ± 95% CI (B, C). Organelle data are presented per cell ($n$ = 42/group) and as the average of the 14 cells/animal (D–F) or per mitochondrion for each animal (G). Data were analyzed by unpaired Student's $t$-test with Welch's correction or Mann-Whitney test, based on normality. *$p < 0.05$ & ****$p < 0.0001$.

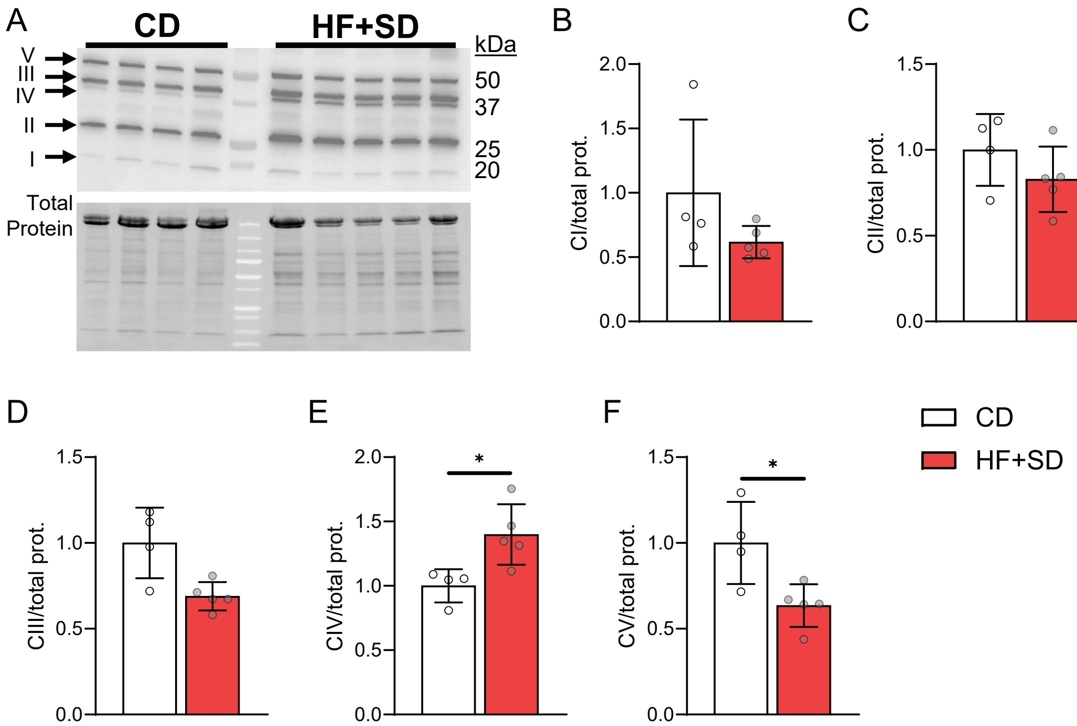


**Supplemental Figure 7. Overnutrition modulates expression of the electron transport chain complexes.** Male mice were given the HF+SD or CD for 6 weeks. Western blots of thyroid homogenates were performed (**A**) and quantitated for complex I (**B**), complex II (**C**), complex III (**D**), complex IV (**E**), and complex V (**F**). MW markers in total protein blot span 10-250 kDa. Data were analyzed by unpaired Student's *t*-test with Welch's correction (B–D, F) or Mann-Whitney test (E). *$p < 0.05$.

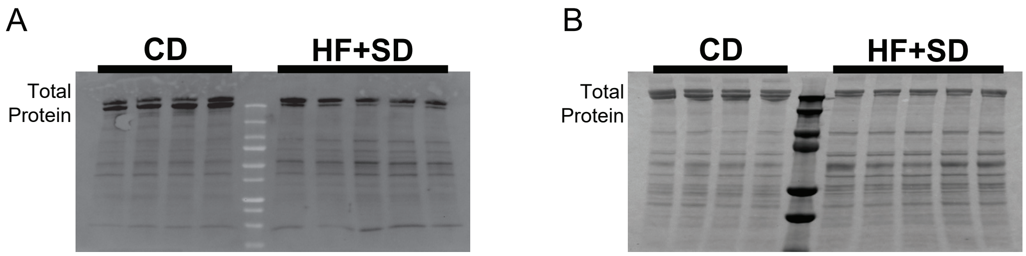


**Supplemental Figure 8. Total protein blots were used to normalize ER stress and TG Western blots.** Total protein per lane was quantitated from the blots later used for BiP and CHOP (**A**) and TG (**B**) expression. MW markers span 10-250 kDa (A) or 25-250 kDa (B). The region with a bubble was consistently excluded across all lanes of the blot.

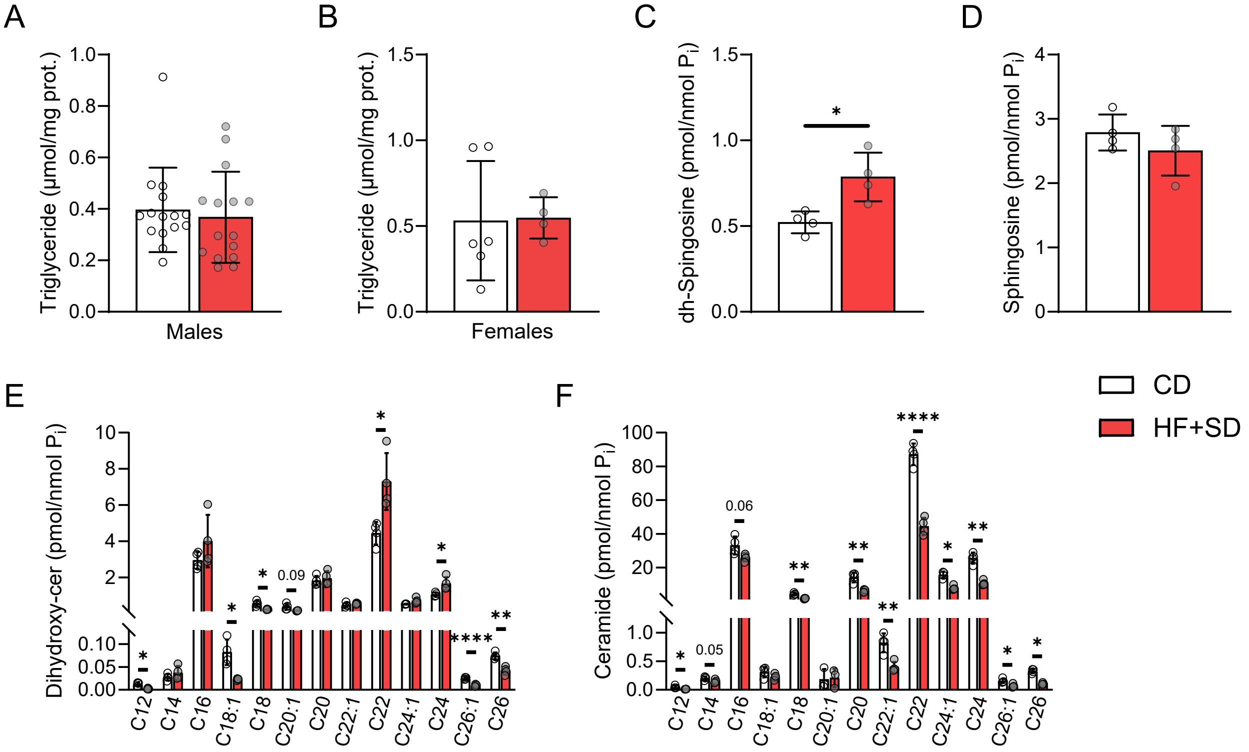


**Supplemental Figure 9. Overnutrition does not result in thyroidal triglyceride or ceramide accumulation.** Mice were placed on the HF+SD or CD for 6 weeks, in the case of the male mice, or 7 weeks, in the case of the female mice. Triglyceride concentration in thyroid homogenates was measured by enzymatic assay for male (**A**) and female (**B**) mice and normalized to the thyroidal protein. Lipidomic analysis of male thyroid tissue was performed for dihydroxy-sphingosine (**C**), sphingosine (**D**), dihydroxy-ceramide species (**E**), and ceramide species (**F**) and normalized to inorganic phosphate concentration. $n$ = 4/group (C–F). Data were analyzed per species by unpaired Student's $t$-test with Welch's correction or Mann-Whitney test, based on normality. *$p < 0.05$, **$p < 0.01$, & ****$p < 0.0001$.

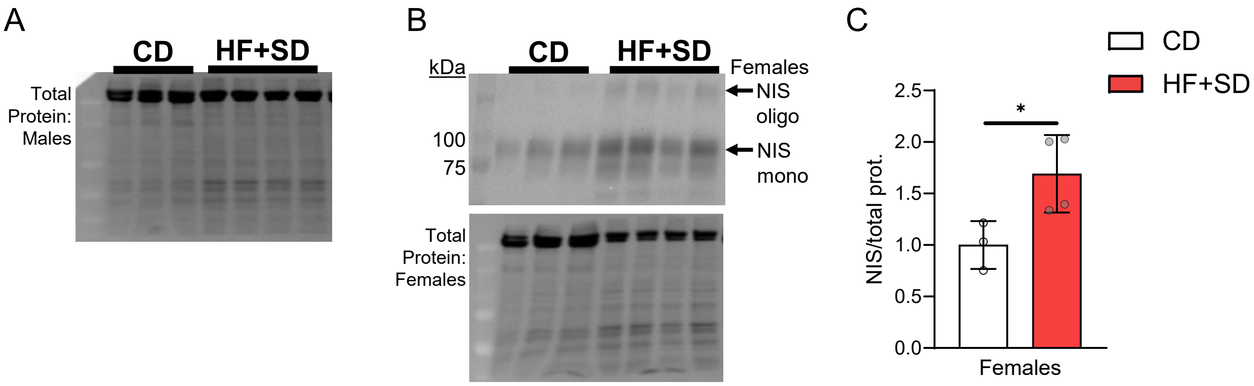


**Supplemental Figure 10. NIS expression is responsive to elevated serum TSH levels.** Total protein per lane was quantitated from the blot later used for NIS expression in male mice (**A**). Female mice were given the HF+SD or CD for 7 weeks. Western blot of thyroid homogenates was performed for NIS (**B**). The lower arrow indicates fully glycosylated monomeric NIS. The upper arrow indicates oligomeric NIS, and the other bands correspond to partially glycosylated NIS. NIS expression in female mice was quantitated (**C**). MW markers in total protein blots span 37-250 kDa. Data were analyzed by unpaired Student's *t*-test with Welch's correction. *$p < 0.05$.

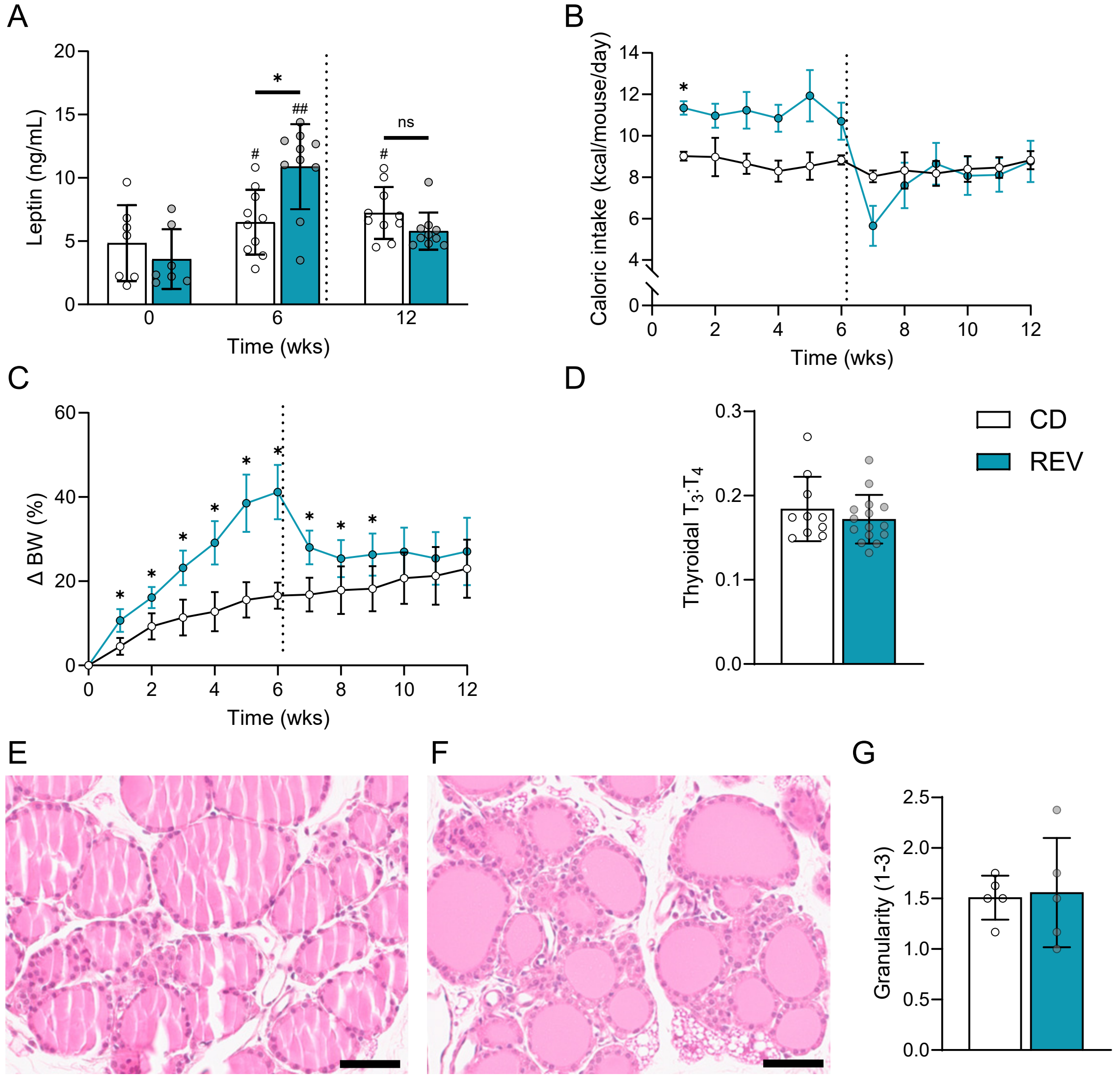


**Supplemental Figure 11. REV mice are weight- and calorie-matched to controls and exhibit normal leptin, thyroid output, and thyroid histology.** Male mice were placed on the HF+SD for 6 weeks and then switched to the CD for 6 weeks (REV). Plasma and sera were analyzed by immunoassay for leptin (**A**). Caloric intake was monitored weekly (**B**). BW was also measured weekly and cumulative percent BW change from baseline was calculated per mouse (**C**). Some thyroids were fully proteolyzed, liberated $T_4$ and $T_3$ were measured by immunoassay, and thyroidal $T_3$:$T_4$ was then calculated (**D**). Some thyroids were fixed, and representative H&E images from a CD mouse (**E**) and a REV mouse (**F**) are shown. H&E images were blindly scored for extent of granularity (**G**). Scale bars = 50 µm. Data are representative of two separate cohorts of mice: this cohort $n$ = 15/group in 3 cages/group (B, C). Data are results from two separate cohorts of mice: combined CD $n$ = 10, HF+SD $n$ = 15 (D). Dotted line represents switch from HF+SD to CD for REV mice. Data were analyzed by 2-way ANOVA with repeated measures (A–C), Mann-Whitney test (D), or unpaired Student's $t$-test with Welch's correction (G). *$p < 0.05$ vs. controls within that timepoint. #$p < 0.05$ & ##$p < 0.01$ vs. week 0 within that diet group.

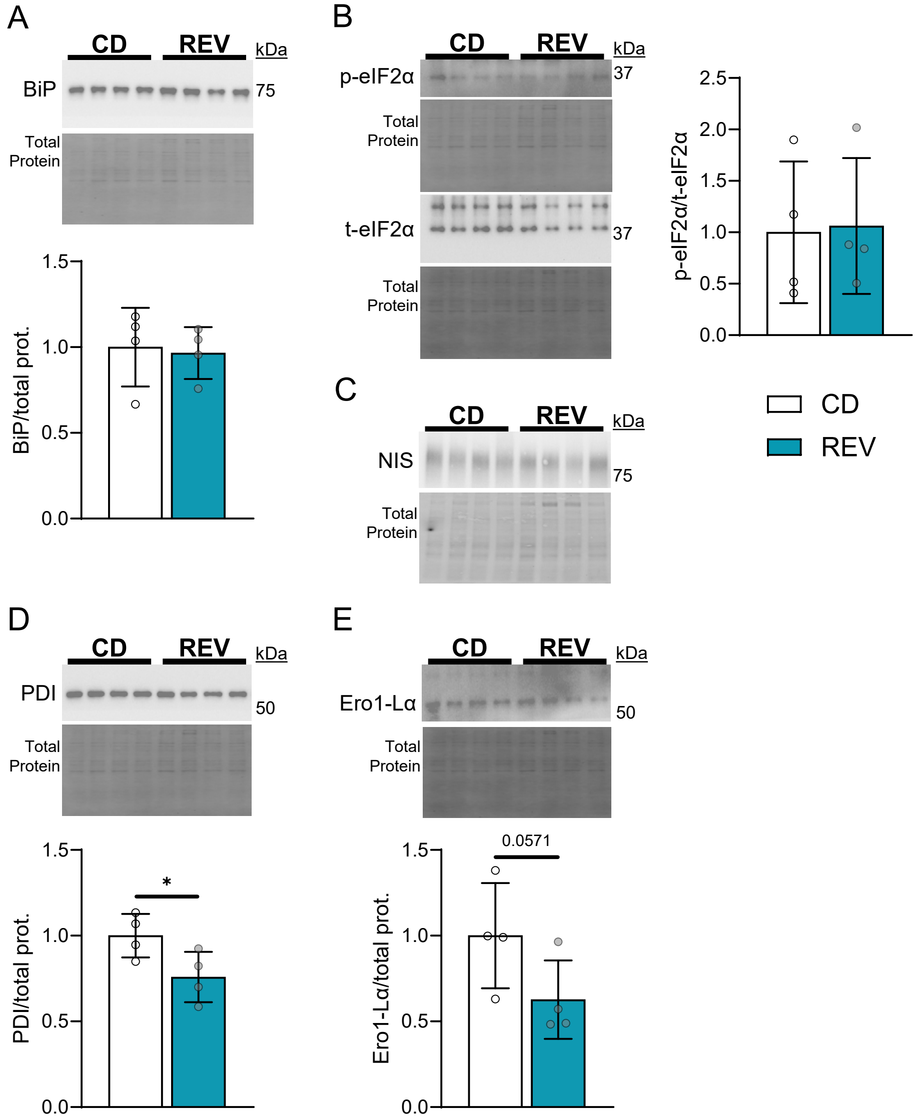


**Supplemental Figure 12. Weight loss reverses thyroidal ER stress and expression of NIS and TG-related proteins.** Male mice were placed on the HF+SD for 6 weeks and then switched to the CD for 6 weeks (REV). Western blots of thyroid homogenates were performed and quantitated for ER stress markers BiP (**A**) and p-eIF2α/t-eIF2α (**B**). Western blot was performed for NIS but not quantitated due to bubbles (**C**). Western blots were also performed and quantitated for TG's folding chaperones PDI (**D**) and Ero1-Lα (**E**). Total protein images for p-eIF2α and PDI are identical (B, D), as are those for t-eIF2α and Ero1-Lα (B, E), because the same blot was cut and used to probe for each pair of targets. MW markers in total protein blots span 15-250 kDa, except the NIS total protein blot, which spans 25-150 kDa. Data were analyzed by unpaired Student's *t*-test with Welch's correction (A, B, D) or Mann-Whitney test (E). *$p < 0.05$.

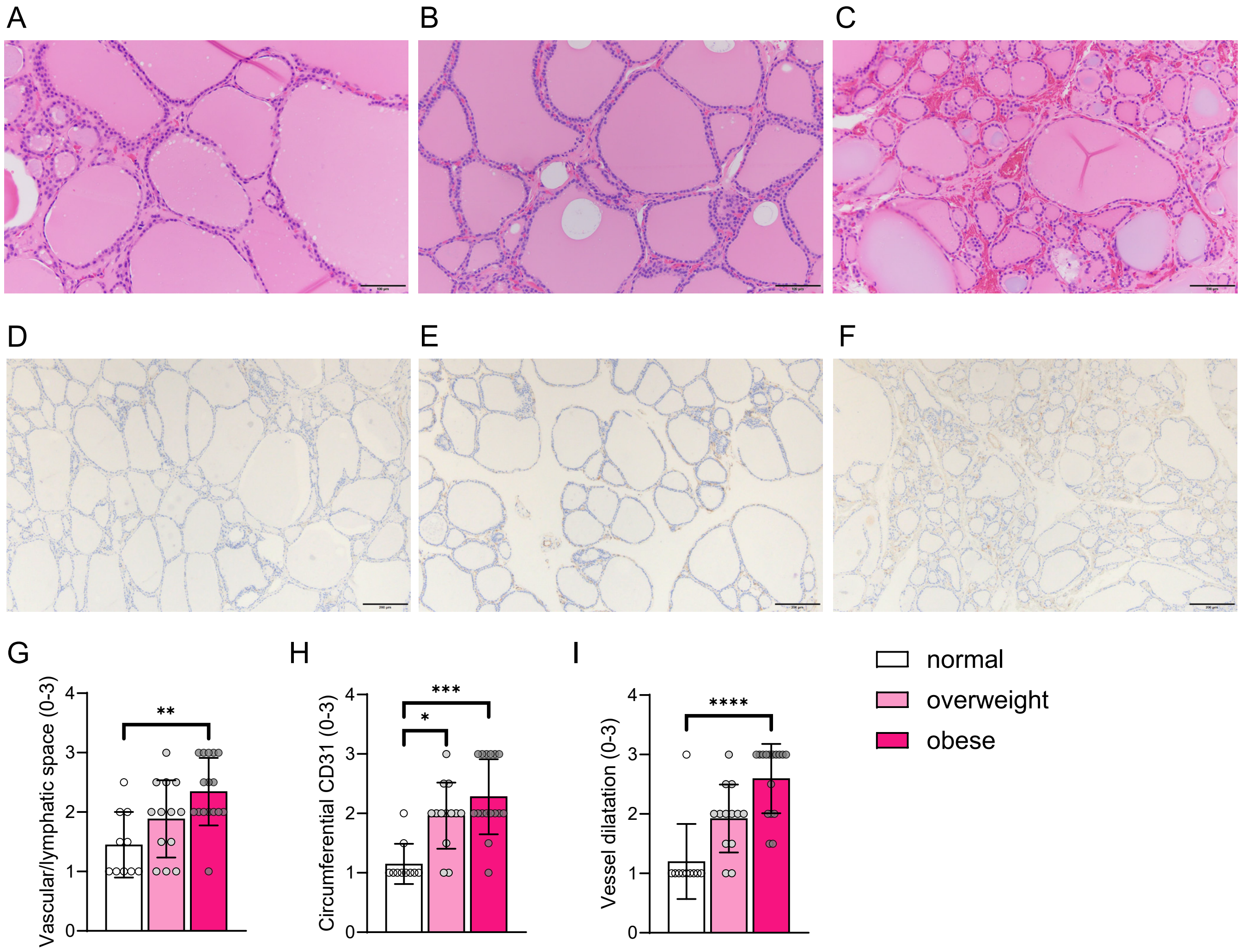


**Supplemental Figure 13. Thyroidal vascularization increases with increasing BMI in humans.** H&E-stained sections of thyroid tissue from patients with MNG were obtained. Representative images from normal (**A**), overweight (**B**), and obese (**C**) subjects are shown. Fixed thyroid sections were stained for vascular marker CD31, and representative images from normal (**D**), overweight (**E**), and obese (**F**) subjects are shown. H&E images were blindly scored for the extent of vascular/lymphatic space (**G**). CD31 images were blindly scored for extent of circumferential staining around the follicles (**H**) and dilatation of the follicular microcapillaries (**I**). Scale bars = 100 μm (A–C) or 200 μm (D–F). Normal BMI = 18.5-24.9, $n$ = 10; overweight BMI = 25-29.9, $n$ = 13; obese BMI ≥ 30, $n$ = 16. Data were analyzed by Kruskal-Wallis test with Dunn's multiple comparison test. *$p < 0.05$, **$p < 0.01$, ***$p < 0.001$, & ****$p < 0.0001$.

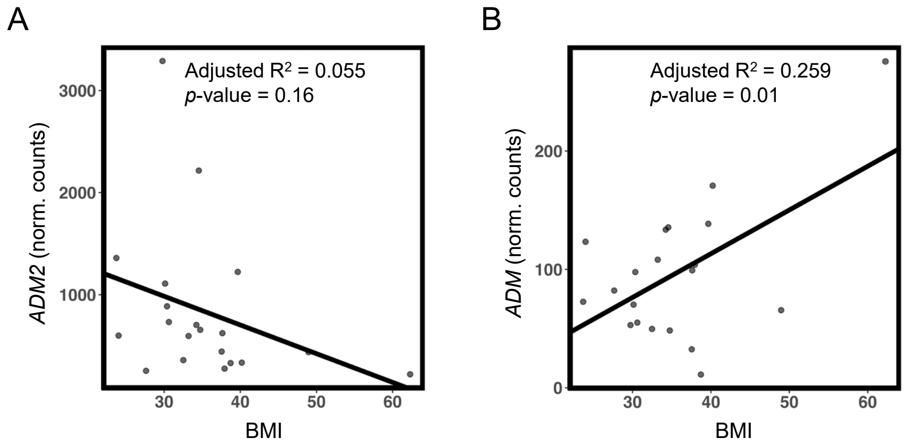


**Supplemental Figure 14. BMI is positively correlated with thyroidal *ADM*.** Bulk RNA sequencing of FFPE thyroid tissue that had been surgically resected from patients with MNG was performed. Gene counts were normalized by DESeq2. Pearson's correlation tests were performed between each patient's BMI and their *ADM2* (**A**) and *ADM* (**B**) expression. $n$ = 20.

**Supplemental Table 1. Top 10 most up- and downregulated gene sets using GO annotations.**

| GO gene set | NES |
|---|---|
| GOBP attachment of spindle microtubules to kinetochore | 2.41 |
| GOBP renal system vasculature development | 2.31 |
| GOBP glomerulus development | 2.26 |
| GOBP kinetochore organization | 2.25 |
| GOBP integrin mediated signaling pathway | 2.25 |
| GOMF microtubule motor activity | 2.24 |
| GOCC condensed chromosome centromeric region | 2.23 |
| GOBP regulation of endothelial cell differentiation | 2.23 |
| GOBP mitotic sister chromatid segregation | 2.23 |
| GOBP protein localization to condensed chromosome | 2.22 |
| GOBP mitochondrial respiratory chain complex assembly | -2.83 |
| GOBP mitochondrial electron transport NADH to ubiquinone | -2.84 |
| HP abnormality of the mitochondrion | -2.86 |
| HP abnormal activity of mitochondrial respiratory chain | -2.87 |
| GOCC NADH dehydrogenase complex | -2.88 |
| GOCC contractile fiber | -2.90 |
| GOCC ribosomal subunit | -2.90 |
| GOMF structural constituent of ribosome | -2.92 |
| GOCC inner mitochondrial membrane protein complex | -3.10 |
| GOCC mitochondrial protein containing complex | -3.20 |

Thyroids were processed for RNA sequencing, and GSEA was performed using GO annotations. Gene sets were ranked by normalized enrichment score (NES) of the HF+SD group vs. controls. CD $n$ = 3, HF+SD $n$ = 4. All adjusted $p$-values < 0.0002.

**Supplemental Table 2. Top 10 most downregulated gene sets using Reactome annotations.**

| **Reactome gene set – 3 weeks** | **NES** |
|---|---|
| Regulation of expression of slits and robos | -2.12 |
| Selenoamino acid metabolism | -2.35 |
| rRNA processing | -2.37 |
| Activation of the mRNA upon binding of the cap binding complex and eIFs and subsequent binding to 43S | -2.38 |
| SRP-dependent co-translational protein targeting to membrane | -2.41 |
| Influenza infection | -2.44 |
| Nonsense-mediated decay | -2.47 |
| Eukaryotic translation initiation | -2.48 |
| Response of eIF2AK4 GCN2 to amino acid deficiency | -2.51 |
| Eukaryotic translation elongation | -2.53 |

Male mice were given the HF+SD or CD for 3 weeks, and their thyroids were processed for RNA sequencing. GSEA was conducted using Reactome annotations. Gene sets were ranked by NES of the HF+SD group vs. controls. $n = 4$/group. All adjusted $p$-values < 0.00001.

**Supplemental Table 3. Spearman correlations between BMI and thyroidal vascularization scores.**

| | BMI vs. vascular/ lymphatic score | BMI vs. CD31 circumference score | BMI vs. CD31 vessel dilatation score |
|---|---|---|---|
| Spearman r | 0.4804 | 0.5428 | 0.5933 |
| *p*-value | **0.0020 | ***0.0004 | ****<0.0001 |

H&E- and CD31-stained thyroid sections from patients with MNG were blindly scored for extent of vascular/lymphatic space, circumferential staining around the follicles, and dilatation of the follicular microcapillaries. Spearman correlation tests were performed between each patient's BMI and each of their 3 histological scores. Normal BMI = 18.5-24.9, $n$ = 10; overweight BMI = 25-29.9, $n$ = 13; obese BMI ≥ 30, $n$ = 16.

**Supplemental Table 4. Primary antibodies.**

| Primary Antibody | Catalog # |
|---|---|
| BiP | CST #3177 |
| p-eIF2α | CST #3398 |
| eIF2α | CST #5324 |
| CHOP | CST #2895 |
| TG | SC #365997 |
| PDI | CST #3501 |
| Ero1-Lα | CST #3264 |
| vinculin | CST #13901 |
| NIS (α-rat) | Generated and affinity purified in the Carrasco lab (1) |
| β-actin | SC #47778 |
| D2 | Abcam #77779 |
| caveolin-1 | CST #3238 |
| OxPhos complexes | Invitrogen #45-8099 |
